# A common structure in recurrent networks supports neural sequence generation locally and in downstream neurons

**DOI:** 10.64898/2026.09.02.748960

**Authors:** Lea Marie Braun, Mathias Karsrud Nordal, Silje-Neneva Hanssen Rambø, Claudia Clopath, Soledad Gonzalo Cogno

## Abstract

Neural sequences, characterized by neurons or groups of neurons that fire one after the other, have been observed in multiple brain regions, across species, and are known to underlie a diversity of brain functions. To flexibly support behaviour and cognition, neural sequences exhibit much variability in properties like their temporal width, baseline, and peak firing rate. Despite this variability and the central role that sequences play in supporting brain function, a framework that explains how flexible sequences are generated and dynamically maintained within a circuit is still missing. Here we go beyond traditional approaches that investigate a one-to-one relationship between network connectivity and specific sequential dynamics. Instead, we train recurrent neural network models to generate a repertoire of sequential dynamics and characterize the obtained connectivity matrices. We found that different connectivity matrices can generate the same neural sequence, yet all connectivity matrices that generate a specific sequence share a common connectivity profile, defined here as the average weight between pairs of neurons as a function of their distance in the sequence ordering. It is the connectivity profile, as opposed to the connectivity matrix, that serves as a fingerprint of the sequential dynamics and shapes the network response to perturbations of the neural activity. Our model predictions were consistent with results obtained from experimental data recorded across brain regions and across species. Finally, we demonstrated that neural sequences can facilitate and constrain the formation of a large repertoire of sequences in downstream brain regions, with the potential of acting as scaffolds for a wide range of computations. Altogether, our results explain how network connectivity can generate a diversity of neural sequences across circuits and how those sequences can be flexibly adapted. Our framework reveals sequences as a common algorithm to support brain function across brain regions and species.

## INTRODUCTION

Sequences of neural activity are characterized by neurons or groups of neurons that fire one after the other. Neural sequences have been reported in many brain regions [1] such as the hippocampus [2-7] and the visual cortex [8-10], and also across species, for example in rodents [3, 11, 12], bats [13], zebrafish [14], turtle [15] and humans [16, 17]. Moreover, neural sequences are known to underlie multiple brain functions, including working memory [7, 18], songbird learning [19, 20], and the representation of time [5, 7, 11, 13]. The ubiquity of sequential activity across brain regions, species, and brain functions suggests that neural sequences constitute a universal pattern of neural activity that can be dynamically adapted to support flexible computations.

Decades of theoretical work have demonstrated that neural sequences can be generated in a neural network with predefined connectivity, such as feed-forward networks [21-23], synfire chains [24-26], or continuous attractor networks [27-30]. Sequences can also emerge in the absence of prewired connectivity, due to plasticity mechanisms [31, 32] or training algorithms [33]. These existing models have primarily focused on how a given network architecture supports one specific neural sequence. However, a theoretical framework that explains how a diversity of sequences can be generated, flexibly modified, and dynamically maintained in neural circuits is still missing.

In this study, we constructed recurrent neural network models that were trained to generate sequences of activity with a wide range of properties. We characterized the connectivity matrices that gave rise to those sequences, and their associated connectivity profiles, defined as the average weight between pairs of neurons as a function of their distance in the sequence ordering. The model predictions were consistent with connectivity inferred from experimental data [12, 13]. By pushing the dynamics out of the sequential regime through local and global perturbations of the network activity, we investigated how the network response was controlled by features of the connectivity profile. Finally, we found that neural sequences can facilitate the formation of a large repertoire of sequences in downstream brain regions. We demonstrate that those sequences are constrained by the properties of the upstream sequence and its corresponding connectivity profile.

## RESULTS

### The multiplicity of connectivity matrices that generate the same neural sequences share a common connectivity profile

We started by characterizing the repertoire of connectivity matrices that give rise to sequences of neural activity. To do this, we trained the weights of rate-based recurrent neural networks (RNNs) to generate sequences with different properties and periodic boundary conditions [33]. We first simulated the target firing rate of each unit in the RNN as a Gaussian kernel that repeated over time and was specified by three parameters: (i) its *amplitude*, corresponding to the difference between the cell’s peak firing rate and its baseline firing rate; (ii) its *offset*, corresponding its baseline firing rate; and (iii) its standard deviation, or *width*, corresponding to the time interval during which it exhibits sustained activity (Figure 1A). In each simulation, these three parameters were fixed at the same value for all network units. To generate sequential activity at the population level in this network of homogeneous neurons, we shifted the activity of each unit in time such that the time points at which the units’ activity peaked uniformly covered the interval from 0 to 8.5 s. Because we established periodic boundary conditions in the temporal domain, the ‘target sequences’ were periodic (Figure 1B, Supp. 1A–D). To reduce the computational time, we used randomly chosen snippets of the target sequences as targets to FORCE train the weights of the RNN [33-36] (Figure 1C). We repeated the training procedure 50 times. In each of the 50 ‘training realisations’ we: (a) randomly initialized the connectivity matrix, (b) used as targets to FORCE train the weights a randomly chosen snippet of the target sequences (Figure 1B) – note that each snippet, of 17 s of duration, contained identical sequences up to a shift in their phase, and (c) quantified the training performance by computing the Pearson correlation between the targets (snippet of the target sequences) and the activity of the trained network. If the training was successful, the Pearson correlation was close to one and the network generated the desired sequential dynamics. Finally, we computed the mean connectivity matrix over the connectivity matrices obtained across training realisations. We also calculated the corresponding ‘connectivity profile’, defined here as the average weight between pairs of neurons as a function of their distance in the sequence ordering (Figure 1C bottom).

**Figure 1:**
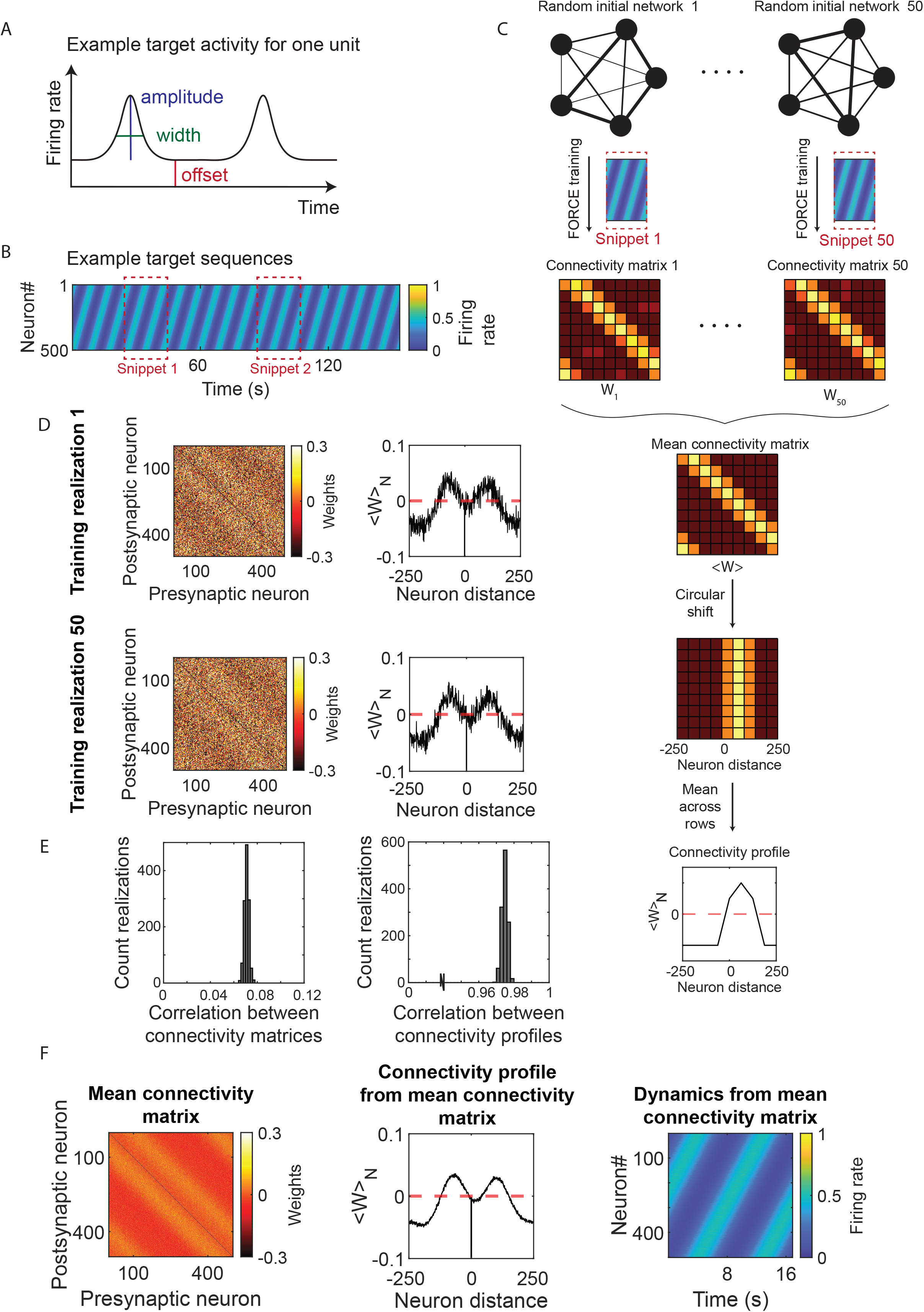
Different connectivity matrices can generate the same sequential dynamics, and all those matrices share the same connectivity profile. **A**, Schematics of simulated target activity for one example unit in the network. We built a Gaussian kernel that repeated over time and which was characterized by three parameters: its amplitude (blue), width (green), and offset (red). **B**, Example target sequences obtained by shifting the units’ target activity in time. Each row indicates the target activity of one of the 501 units in the network. Cold colours indicate low firing rate, warm colours high firing rate. The firing rate, by construction, is bounded between 0 and 1. Snippets of the sequential activity, marked in red, were randomly chosen. Each sequence was 8.5 s long. Throughout this figure we used: Amplitude = 0.5, Width = 1.7 s, Offset = 0. **C**, Flow diagram of the procedure for calculating the mean connectivity matrix and its corresponding connectivity profile. From top to bottom: First, we built 50 RNNs that were fully and randomly connected. Next, we trained each RNN separately, yielding a total of 50 training realisations, depicted in the diagram from left to right. In each of the 50 training realisations, we chose a random 17 s snippet from the target sequences (see panel B) and used it to FORCE train the recurrent weights of the RNN. Each training realisation yielded a connectivity matrix, depicted as a heat map in shades of red. Dark red indicates negative weights, yellow indicates positive weights. Presynaptic neurons are shown along the x axis, and postsynaptic neurons along the y axis. We next applied a circular shift to each row of the resulting connectivity matrix such that the weights of the self-connections were centered in the middle column. To compute the connectivity profile, we calculated the mean across rows. The red dashed line in the bottom panel is positioned at zero weights. **D**, Left: Connectivity matrices obtained from training realisation 1 (top) and training realisation 50 (bottom). Correlation value between both matrices: 0.07. Right: Connectivity profiles calculated on the connectivity matrices on the left. Correlation value between both connectivity profiles: 0.98. Symbols and colours as in panel C. **E**, Left: Distribution of correlation values calculated between pairs of connectivity matrices from the total of 50 training realisations. Right: Distribution of correlation values calculated between pairs of connectivity profiles obtained in the 50 training realisations. The connectivity matrices differ across training realisations, but the connectivity profiles are very similar. **F**, Left: Mean connectivity matrix calculated across all 50 training realisations. Middle: Connectivity profile calculated on the mean connectivity matrix. Right: Network dynamics generated with the mean connectivity, which resembles the target sequences (right). Symbols and colours as in panel B, C.

The training was successful for a wide range of amplitude, offset and width values (Supp. 1E, F). For each combination of those three parameters, which defined sequences with those specific properties (Supp. 1A-D), the 50 training realisations produced different connectivity matrices (Figure 1D left, E left, Supp. 1G). However, despite the differences in connectivity matrices, the corresponding connectivity profiles were remarkably similar (Figure 1D right, E right). Moreover, they closely matched the connectivity profile computed on the mean connectivity matrix across training realisations (Figure 1F). Furthermore, the connectivity profiles remained similar when the RNNs had sparse connectivity (Supp. 2). These results show that while a multiplicity of connectivity matrices can generate the same target sequences, those matrices share a unique connectivity profile whose features are constrained by the sequential dynamics. We demonstrate this result analytically by showing that each connectivity matrix can be decomposed into a structured and an unstructured component. The former is the same across matrices, contains information about the target sequences, is represented in the connectivity profile, and its rank is determined by the number of dominant Fourier components of the target sequences. The latter varies across training realisations (see Supp. Material).

### Changes in the connectivity profile map changes in the properties of the neural sequences

We next investigated the relationship between features of the connectivity profile and properties of the target sequences. First, we observed that sequences emerged from an asymmetric connectivity profile (Figure 2A left). This asymmetry supports the temporal progression of the sequence by enabling the propagation of localized activity, or ‘bump of activity’, throughout the network. The asymmetry increased with increasing offset values (Figure 2A, Supp. 3A). Furthermore, when we increased the width of the sequences the range of the connectivity profile, defined as the difference between its maximum and its minimum, decreased (Figure 2B, Supp. 3B). Finally, increasing the sequence amplitude preserved the asymmetry in the connectivity profile while it increased its range (Figure 2C). Interestingly, an oscillation of increasing frequency appeared (Figure 2C, Supp. 3C). These results predict which features the connectivity profile should have in order to support specific target sequences. Moreover, they show that smooth changes in the neural sequences are supported by gradual changes in the connectivity profile (Supp. 4, Supp. Material).

**Figure 2:**
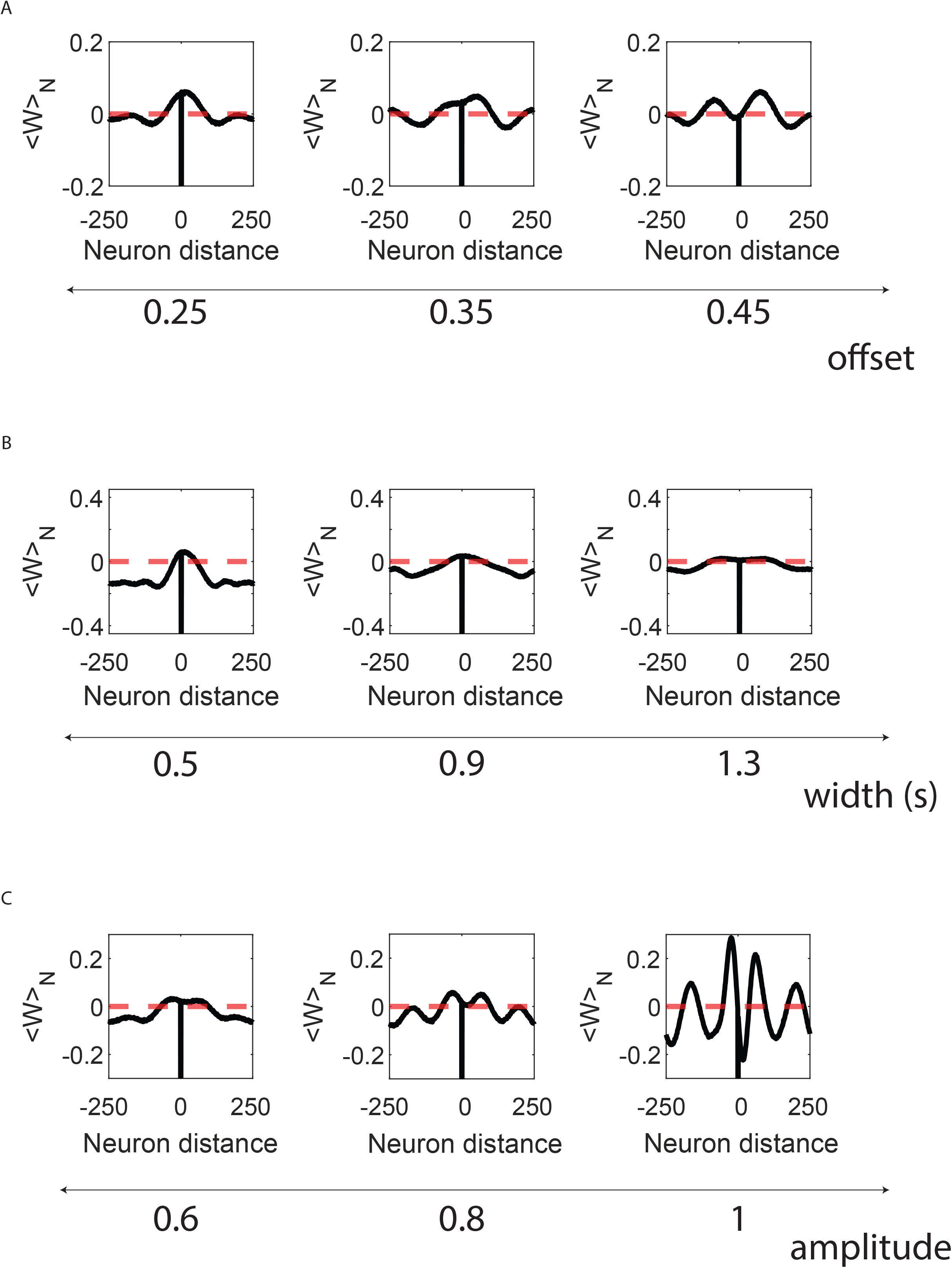
Changes in the connectivity profile smoothly track changes in the properties of the target sequences. **A**, From left to right: Connectivity profiles for increasing offset values. The profile becomes increasingly asymmetric. Amplitude = 0.5, Width = 1.0 s. **B**, Same as panel A, but for increasing width values. The ‘range’ of the profile, defined as its maximum minus its minimum, decreases. Amplitude = 0.5, Offset = 0. **C**, Same as panel A, but for increasing amplitude values. An oscillation of increasing frequency appears. Width = 1.0 s, Offset = 0. Symbols and colours as in Figure 1C.

Previous work has shown that RNN dynamics are shaped not only by global features of the connectivity matrix, but also by higher order statistics [37-42]. We next quantified the presence of connectivity motifs in the connectivity matrices that emerge from training (Supp. 5A). We found that increasing the target sequences’ offset, width, or amplitude led to a higher abundance of reciprocal connectivity motifs (Supp. 5B). Interestingly, we did not find any divergent, convergent nor chain connectivity motif, suggesting that these motifs, and in particular chain motifs, are not strictly necessary for supporting sequences of activity (Supp. 5B). These results extend the connection between connectivity profiles and target sequences by demonstrating that changes in the target sequences are accompanied by gradual changes in the connectivity motifs.

### Connectivity profiles inferred from experimental data match model predictions

For target sequences with specific values of amplitude, width, and offset, our model predicts its associated connectivity profile. We next tested the model predictions on experimental data. We focused on two different and complementary datasets. In one dataset two bats, a demonstrator bat and an observer bat, were in a room with three landing balls, ‘start’, ‘A’, and ‘B’ [13]. Electrophysiological recordings from the hippocampal CA1 showed that time cells exhibit sequential activity when both bats were hanging motionlessly from one of the landing balls (Figure 3A). In the other dataset, head-fixed mice run at their own pace on a wheel [12]. Calcium imaging recordings from the medial entorhinal cortex (MEC) showed that neural activity was organized into ultraslow periodic sequences of activity (Figure 3B).

**Figure 3:**
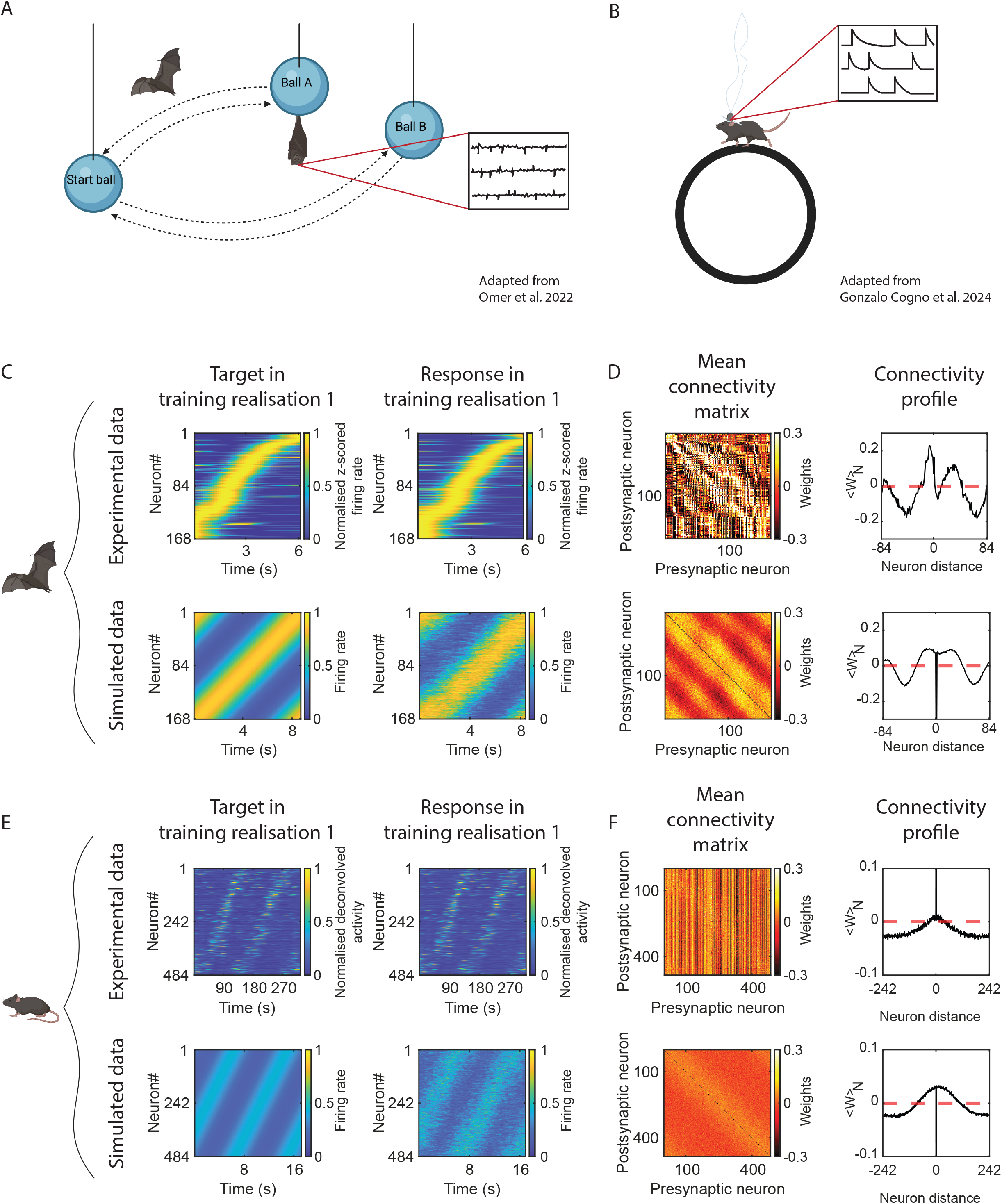
Model predictions agree with connectivity inferred from experimental data. **A**, Schematics of experimental protocol in which electrophysiological recordings from the CA1 of flying bats revealed that neural activity was organized into sequences. Two bats were placed in a room with three landing balls. One bat, the demonstrator, was trained to fly from the start ball to either ball A or B and back. The second bat, the observer, was trained to imitate the behaviour of the demonstrator. We analysed data from the observer bat. We used the neural data collected while the observer bat was hanging still from either ball A or B. Adapted from Omer, Las, and Ulanovsky 2023. Created in BioRender. Gonzalo Cogno, S. (2026) https://BioRender.com/q9qr76c. **B**, Schematics of experimental protocol in which calcium imaging recordings from the medial entorhinal cortex (MEC) of mice revealed that neural activity was organized into sequences. The mice were head fixed and placed on a running wheel in sensory minimized conditions, and without any landmarks or rewards. Adapted from Gonzalo Cogno et al., 2024. **C**, Left: Targets generated with CA1 bat experimental data (top) or simulated data constructed to best approximate the experimental data (bottom). We show the target and the RNN response corresponding to training realisation 1. Parameters of the simulated targets: Amplitude = 0.83, Width = 1.72 s, Offset = 0.05. Right: Response of the RNN after training on the targets depicted on the left. Pearson correlation between target and RNN response: 0.90 (experimental target) and 0.90 (simulated target). Symbols and colours as in Figure 1B. Bat schematics: Created in BioRender. Gonzalo Cogno, S. (2026) https://BioRender.com/481uxqh. **D**, Left: Mean connectivity matrix calculated on networks trained on CA1 bat experimental (top) and simulated (bottom) data. Correlation between the top and bottom connectivity matrices: 0.13. Right: Connectivity profiles corresponding to the matrices on the left. Correlation between connectivity profiles: 0.83. Symbols and colours as in Figure 1C. **E**, Similar to C but for MEC mouse experimental data. Parameters of the simulated targets: Amplitude = 0.29, Width = 1.68 s, Offset = 0.15. Pearson correlation between target and RNN response: 0.95 (experimental target) and 0.69 (simulated target). Mouse schematics: Created in BioRender. Gonzalo Cogno, S. (2026) https://BioRender.com/pdgc94o. **F**, Similar to D but for the RNNs in panel E. Correlation between connectivity matrices: - 0.09. Correlation between connectivity profiles: 0.95.

To infer connectivity from the experimental data we trained two RNNs: one using data from a single mouse recording session, and the other using data pooled across multiple bat recording sessions (see Methods). Each network had as many units as cells in its corresponding dataset (484 in the mouse session, and 168 in the bat dataset). Similarly to what we did for simulated data, we trained the weights of each RNN using the recorded activity of individual neurons as targets, obtained the corresponding connectivity matrices and calculated the connectivity profiles. To compare the connectivity between experimental and simulated data, we first fit a Gaussian kernel to each experimental dataset (Supp. 6) and used the fitted parameters to generate synthetic sequences. Next, we trained a new set of RNNs on these synthetic data and calculated the corresponding connectivity profiles. The RNNs learned all sets of experimental (Figure 3C top, E top) and simulated (Figure 3C bottom, E bottom) targets. Even though the synthetic data was constructed to best approximate the experimental data, the resulting connectivity matrices differed (Fig 3D left, F left). While we did not enforce the Dale’s law while training, the connectivity matrices obtained from experimental data exhibited a band-like structure, indicative of synaptic weights that arise from distinct excitatory and inhibitory neural populations (Figure 3D left top, F left top). We did not find such structure in the connectivity matrices obtained from simulated data (Figure 3D left bottom, F left bottom). Moreover, since the connectivity matrices between experimental and simulated data differed, so did the connectivity motifs (Supp. 7). Differences between the experimental and simulated data may underlie the differences we observed in the connectivity matrices. Yet, the dissimilarity across matrices aligns with our previous findings showing that different architectures can generate the same sequential dynamics. As expected, however, the connectivity profiles were very similar (Figure 3 D right, F right). All together, these results demonstrate that the connectivity profiles obtained from experimental data match those of simulated data. In addition, we further show that a multiplicity of connectivity matrices is consistent with one given connectivity profile. We conclude that it is the average connectivity across neurons that needs to hold in order to support specific dynamics.

### Sequences of activity reset in the presence of perturbations, and those resets are controlled by the connectivity profile

Animals can sense stimuli or engage in behaviours that could disrupt the progression of ongoing neural sequences. To investigate the robustness of sequential dynamics and the link between this robustness and network connectivity, we incorporated perturbations into our simulations. We studied two different types of perturbations: (i) ‘structured’, in which the activity of an ensemble of neighbouring neurons in the sequence is perturbed, and (ii) ‘unstructured’, in which the activity of randomly selected neurons is perturbed.

We begin by describing the effects of structured perturbations. First, we divided neurons into non-overlapping ensembles such that neighbouring cells in the sequence ordering belonged to the same ensemble. Second, we perturbed the activity of a single ensemble during a specific time window (Figure 4A). Third, we quantified the network response by correlating the post-perturbation network activity with that of the unperturbed network. We first investigated the effects of silencing the activity of one ensemble (Figure 4B). In a biological context this perturbation could represent a sudden lack of stimulation affecting a subset of cells. For a wide range of properties of the target sequences, and therefore for different connectivity profiles, we found that silencing an ensemble that would have been active in the sequence resets the ongoing sequence to another ensemble (Figure 4C left, C middle left). As the sequence offset or width increases, the ensemble to which the sequence resets is further away from the perturbed ensemble (Supp. 8A, B). Furthermore, as the sequence amplitude increases, the reset coexists with the emergence of transient parallel sequences that vanish quickly over time (Supp. 8C). These observations can be explained in terms of the connectivity profile. The sequence resets to the ensemble composed of the units receiving the largest total input current, which are those with the weakest inhibition. These units may receive negative weights from the silenced neurons, such that when the presynaptic units are silenced, their contribution to the input current of the postsynaptic units is zero and therefore the net postsynaptic input current increases. These postsynaptic units may therefore be the ones located at the minimum of the connectivity profile relative to the units that were silenced (Supp. 8). In the case of increasing amplitude values, the various sequences that are activated after silencing are enabled by the multiple minima in the connectivity profile (Supp. 8C bottom), which gave rise to the formation of multiple bumps of activity. Finally, and as expected, if the silenced ensemble was not engaging in the sequence during the time of the perturbation, the sequence continued uninterruptedly (Supp. 9A). Our results hold for different ensemble sizes (Supp. 9A).

**Figure 4:**
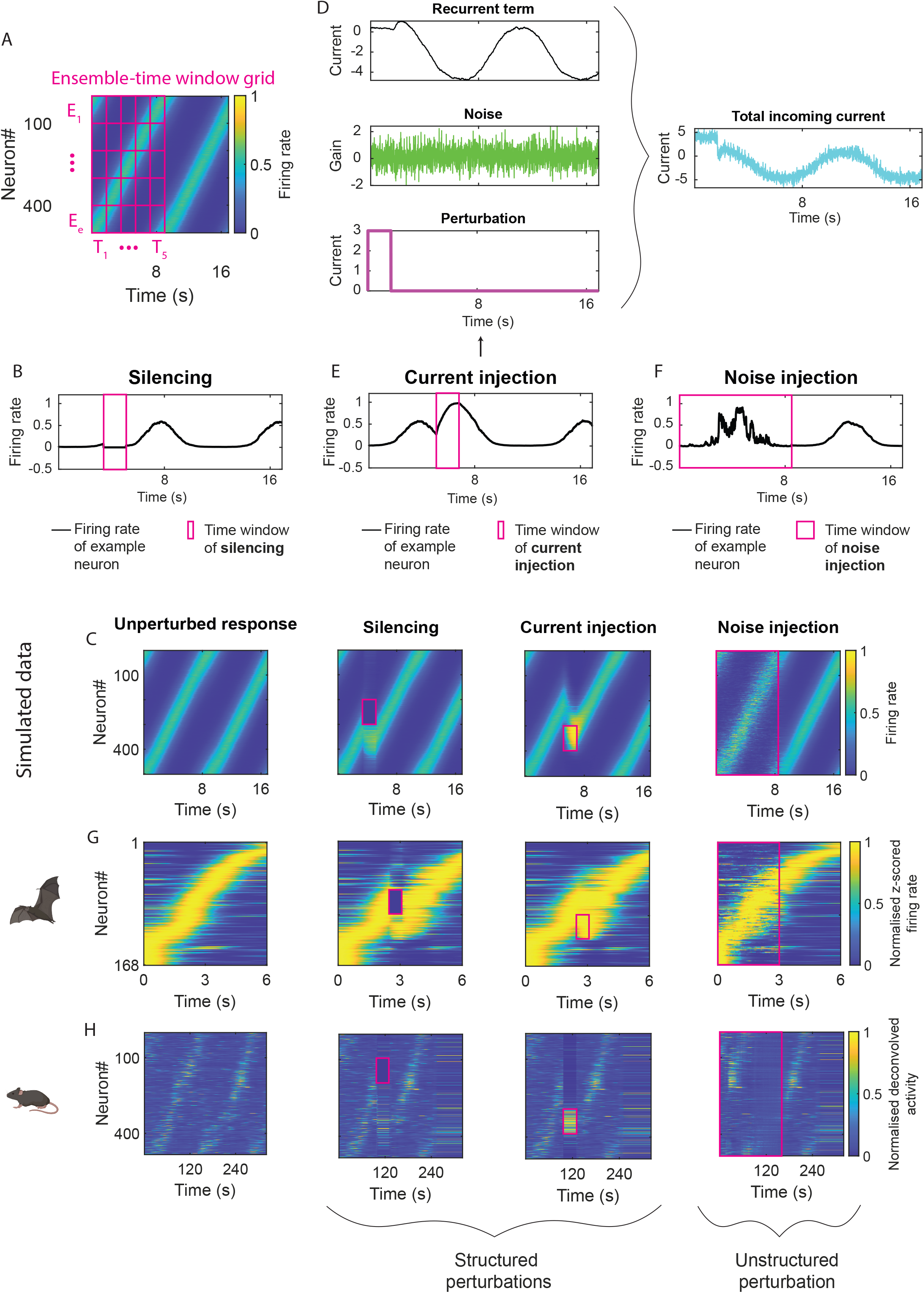
Sequences reset upon structured perturbations, and the reset is controlled by the connectivity profile. **A**, Schematics of the ensemble-time window grid that we used for perturbing the network activity in a structured manner. We split the neurons into *e* equal-sized ensembles of neighboring neurons (E_1_, …, E_e_), and the duration of the first sequence into five equal-sized time windows (T_1_, …, T_5_). The grid is depicted in magenta. The perturbation was applied to one given ensemble, at one specific time window. No perturbations were applied during the second sequence. Properties of the depicted sequence: Amplitude = 0.6, Width = 1 s, Offset = 0. **B**, Impact of the silencing perturbation on the firing rate (black) of an example neuron. The time window during which the perturbation took place is indicated in magenta. The firing rate decreases to 0 during the perturbation. **C**, Left: Unperturbed response of a network consisting of 501 neurons that was trained on synthetic data. Amplitude = 0.6, Width = 1 s, Offset = 0. Duration of a sequence = 8.5 s. Duration of the simulation = 17 s. Middle left: Response of the network on the left following a silencing perturbation applied to the ensemble and time window indicated in magenta. Ensemble size = 100 neurons. Duration of perturbation time window = 1.7 s. Middle Right: Response of the network on the left panel following the injection of additional current into the ensemble, and at the time window, indicated in magenta. Magnitude of the step current = 3. Ensemble size = 100 neurons. Duration of the perturbation time window = 1.7 s. Right: Response of the network on the left panel following the injection of noise into 50% of randomly selected neurons, during the time window of 8.5 s indicated in magenta. Noise gain = 15. **D**, Composition of the total incoming current into one example neuron when additional current was injected. The recurrent term is shown in black, the noise term in green, the injected current in magenta, the sum of all terms in cyan. **E**, Same as B, but in the case of adding additional current. Magnitude of the step current = 3. **F**, Same as E but in the case of adding noise during the first sequence (8.5 s). Noise gain = 15. **G**, Same as in C, but for a network trained on CA1 bat experimental data. The heat maps show the normalized z-scored firing rates of 168 neurons. Middle: Ensemble size = 33 neurons. Duration of the perturbation time window = 0.6 s. Middle right: Magnitude of step current = 3. Right: Percentage of perturbed neurons = 50%, Noise gain = 20. Duration of the perturbation time window = 3 s. **H**, Same as in C, but for a network trained on MEC mouse experimental data. The heat maps show the normalized deconvolved activity of 484 neurons. Middle: Ensemble size = 96 neurons. Duration of the perturbation time window = 32.3 s. Middle right: Magnitude of step current = 3. Right: Percentage of perturbed neurons = 50%, Noise gain = 3. Duration of the perturbation time window = 161.3 s. The sequence resets yet eventually settles into an attractor state. Symbols and colours as in Figure 1B.

We next considered the case in which an ensemble of neurons suddenly receives additional input current. In a biological context, this perturbation may mimic the response of a subset of cells to the onset of a stimulus or environmental feature (Figure 4D, E). We found that the original sequence stops and a new sequence starts from the perturbed ensemble (Figure 4C middle right). The new sequence has the same properties as the original sequence because the connectivity profile remained unchanged. While our findings are independent from the ensemble size, the effect was only present for high enough values of injected current (Supp. 9B, C). Finally, we focused on unstructured perturbations by injecting additional noise into a subset of randomly selected neurons (Figure 4F). We found that the sequential dynamics persisted for a wide range of percentages of perturbed neurons and noise gains (Figure 4C right, Supp. 9D, E).

Finally, we tested the predictions of the model on experimental data. We applied the same perturbations as above to the response of RNNs that were trained to generate experimental recordings from the bat CA1, or mouse MEC (Figure 4G, H). When the network was trained on bat data, it exhibited responses to the perturbations that were consistent with those predicted by the model (Figure 4G, Supp. 9F-H). The response of the network trained on mouse data also exhibited a reset of the sequential activity following structured perturbations, as predicted by the model, yet the dynamics eventually converged to an attractor state (Figure 4H middle, Supp. 9I-K). The attractor state remained present when we applied unstructured perturbations (Figure 4H right).

### Network connectivity supports, and constrains, flexible sequences in downstream brain regions

Sequences of neural activity can act as scaffolds that facilitate computations [7, 11, 13, 18, 19] or the formation of new patterns of activity in downstream regions [9, 12, 43]. This is especially relevant given that sequential activity has been reported in multiple brain areas [1, 5, 12, 19, 44]. We next investigated how sequences of activity shape sequential dynamics in downstream regions. First, we simulated an ‘upstream network’ that consisted of 501 units and that exhibited sequential activity (Figure 5A). Next, we used the activity in the upstream network as input to a ‘downstream network’, which was composed of 10 units. All upstream neurons were connected to all downstream neurons (Figure 5A). We trained those weights to enforce sequential activity in the downstream network. Finally, we considered one specific sequence in the upstream network (obtained after FORCE training as we did in previous sections - parameters: amplitude = 0.5, width = 1 s, offset = 0.3) and asked whether the downstream circuit could learn sequences with varying properties using the predefined upstream sequence as input (Supp. 10A, Figure 5B).

**Figure 5:**
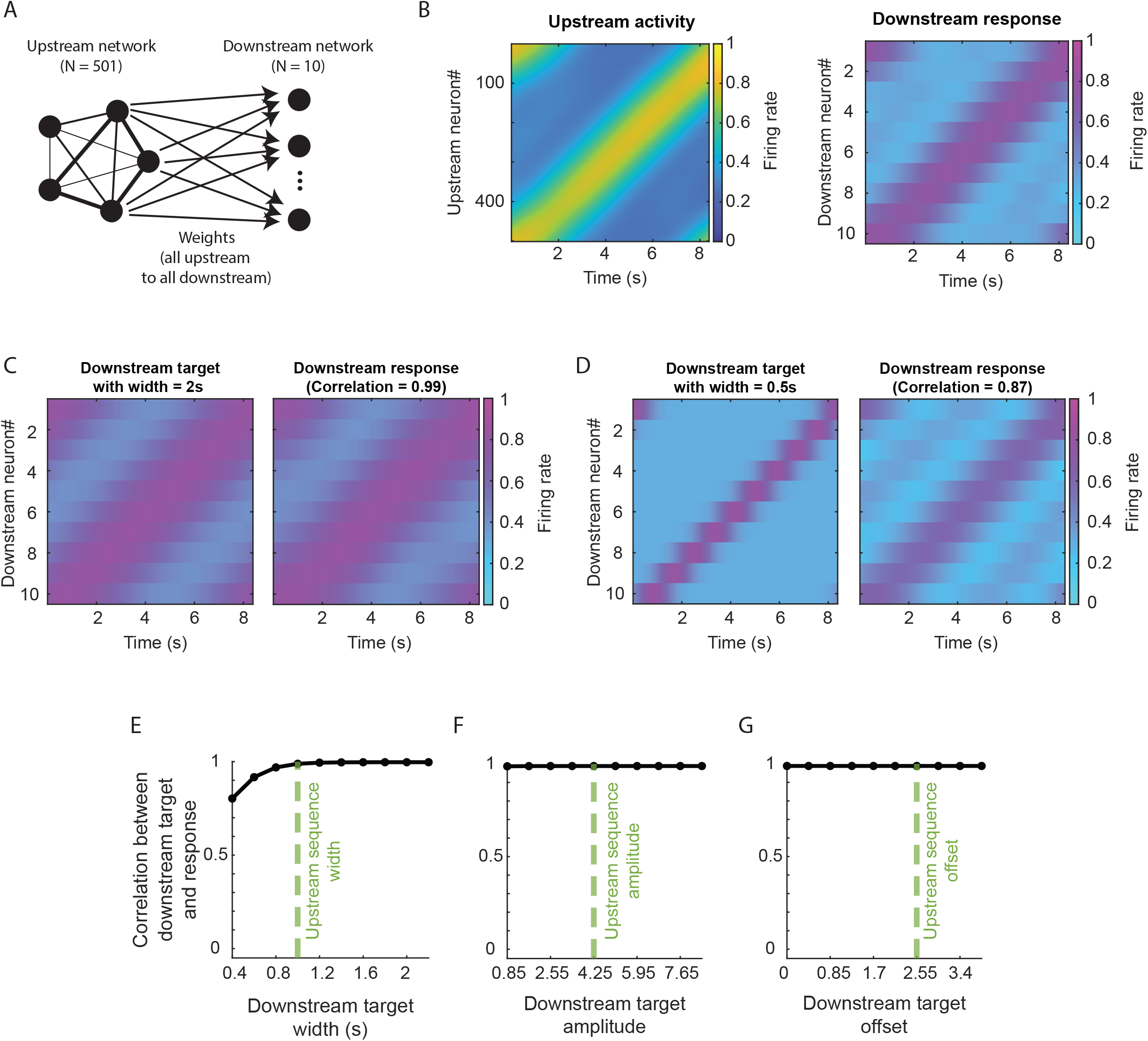
Neural sequences in an upstream area support a diversity of sequences in downstream regions. **A**, Schematics of the two-layers neural network. The ‘upstream network’ is fully connected and composed of 501 neurons. The ‘downstream network’ contains 10 neurons that are not connected to each other. All neurons in the upstream network are connected to all neurons in the downstream network. The upstream network generates sequential activity that serves as input to the downstream network. The weights between the upstream and the downstream neurons are trained so that the downstream network generates sequences with specific properties. **B**, Left: Example upstream network activity that is used as input to the downstream network. Amplitude = 0.5, Width = 1 s, Offset = 0.3. Sequence duration = 8.5 s. Colours as in Figure 1B. Right: Response of the downstream network receiving as input the sequence on the left, and after training. Parameters of the downstream target: Amplitude = 0.5, Width = 1 s, Offset = 0.3. Sequence duration = 8.5 s. Cyan indicates low firing rate and magenta indicates high firing rate. **C**, Left: Downstream target parameters: Amplitude = 0.5, Width = 2 s, Offset = 0.3. The downstream target is wider than the upstream sequence, depicted in B, left. Right: Activity of the downstream network after training. Correlation between downstream target and response: 0.99. Colours as in Figure 5B. **D**, Same as in C but with these downstream target parameters: Amplitude = 0.5, Width = 2 s, Offset = 0.3. The downstream target is narrower than the upstream sequence, depicted in B, left. Correlation between downstream target and response: 0.87. Colours as in Figure 5B. **E**, Correlation between downstream target and network response as a function of different downstream target width values. Amplitude = 0.5, Offset = 0.3. The upstream sequence was kept fixed with parameters: Amplitude = 0.5, Width = 1 s, Offset = 0.3. The width of the upstream sequence is indicated with a green dashed line. The downstream network could not learn targets that had a smaller width than that of the upstream sequence. **F**, Same as E but over different amplitude values of the downstream target. Width = 1 s, Offset = 0.3. The amplitude of the downstream sequence could be different from the amplitude of the upstream sequence and the training was still successful. **G**, Same as E but over different offset values of the downstream target. Amplitude = 0.5, Width = 1 s. The offset of the downstream sequence could be different from the offset of the upstream sequence and the training was still successful.

We found that the training was successful as long as the sequence in the downstream region was as wide as, or wider than, the sequence in the upstream region (Figure 5C-E). In addition, we found that the same upstream sequence could generate a downstream sequence with a wide range of amplitude and offset values (Figure 5F, G, Supp. 10B, C). Next, we investigated the impact of perturbations in the upstream sequence on the generation of sequences in downstream areas. As expected, we found that if the perturbation is such that the activity in the upstream layer is disrupted, then the activity in the downstream layer is disrupted too (Supp. 10D, E). This result suggests that the scaffolding role is constrained to sequences unfolding in an uninterrupted manner. Our results can be understood analytically (Supp. Material): since the downstream network generates a linear readout of the activity in the upstream layer, the downstream sequence will be shaped by the upstream sequence, and therefore by the connectivity profile in the upstream layer. Altogether, these findings indicate that the connectivity profile in a brain region not only determines which sequences can be generated locally but also supports flexible sequence generation in downstream regions.

## DISCUSSION

The brain is known to generate a big repertoire of sequences of neural activity across multiple brain regions, ranging from visual, auditory, and somatosensory areas, to subcortical regions such as the striatum and the hippocampal formation [1, 7, 9, 11-14, 19, 45, 46]. To understand how a variety of sequences can be generated and dynamically modified within and across circuits, we went beyond traditional approaches that focused on the one-to-one mapping between a predefined connectivity matrix and neural sequences with fixed properties [24, 27, 31, 33]. Instead, we trained RNNs to generate neural sequences with a wide range of properties and investigated the connectivity matrices that support those sequences. We found that different connectivity matrices can support the same sequence, yet all those connectivity matrices share a common connectivity profile. In addition, the connectivity profile changes smoothly as properties of the sequences vary. We showed that sequences reset upon perturbations, and that these resets are controlled by the connectivity profile. Finally, we quantified how the connectivity profile can facilitate, and constrain, the formation of a big repertoire of sequences in a downstream area.

In this work, we found that multiple recurrent connectivity matrices can generate the same sequential dynamics. More specifically, we found that the connectivity matrix can be decomposed into a structured component, and an unstructured residual. Importantly, only the structured component is constrained by the target sequences, while the residual can vary without affecting the sequential activity of the network. These findings reveal a many-to-one relationship between recurrent connectivity and sequential dynamics. This degeneracy is not specific to our study but a general feature of recurrent networks [47, 48]. For example, in low-rank RNNs the structured low-dimensional component of the connectivity matrix determines the dynamics, while the unstructured component is unconstrained [49, 50]. Additionally, recent studies show that the structured component can be implemented in different ways and still give rise to equivalent dynamics [51-53]. Together, these results suggest that even though the many-to-one relationship between connectivity and dynamics is a robust feature of RNNs, networks that implement the same dynamics share a common underlying structure. In the case of sequential activity considered here, this shared structure is the connectivity profile, a circulant component embedded within the recurrent connectivity.

Our findings show that the peak firing rate, width, and baseline activity of the sequences can be dynamically modified by adjusting properties of the connectivity profile such as its degree of asymmetry, its oscillatory frequency and its maximum and minimum values. These specific modifications may enable the formation of new sequences on the fly without the need for substantial circuit rewiring. This flexibility may extend to other properties of the sequences too. For example, it has been shown that increasing the weights between excitatory assemblies shortens the duration of a sequence, and increasing the weights from excitatory to inhibitory assemblies extends it [23]. While network connectivity plays a key role in shaping sequential dynamics, other mechanisms in the brain may also contribute to adjusting the properties of sequential activity, such as the magnitude and temporal structure of external input [54] or the brain’s temperature [55].

The degeneracy of the connectivity matrix, together with the generation of flexible sequences that are enabled by gradual changes in the connectivity profile, may facilitate the brain’s ability to adapt the sequential neural dynamics to varying behaviours and cognitive demands. They may also support the coexistence of different dynamics and codes within the same network. For example, sequences may need to quickly change according to the context or choices in a decision-making task [3, 18], or based on external stimuli [7]. Furthermore, much of the information encoded in a network, such as contextual cues or memory traces of past events, should remain stable regardless of ongoing computations supported by the sequences. The degeneracy of the connectivity matrices may have the computational advantage of preserving stable information representations while enabling flexible and behaviour-dependent sequences.

Our results provide testable predictions for connectome datasets. The degeneracy of the connectivity matrix suggests that simply comparing network connectivity with connectome data may be misleading. Instead, it is the structured component of the connectivity matrix, which in this work is captured by the connectivity profile, that should be compared to experimental data. Because our findings are not restricted to any specific brain region or species, our model predictions could potentially be tested on connectome data collected from different animal models, such as rodent [56], fly [57-59] and zebrafish [60]. All these species are known to exhibit sequences of neural activity.

The many-to-one relationship between recurrent connectivity and sequential dynamics may pose challenges when trying to infer connectivity from neural activity data [61-64]. Such inference is generally difficult even with sophisticated methods, because as we show here, the dynamics constrain the underlying connectivity only weakly [63, 64]. However, knowing that the connectivity must contain a low-rank circulant component provides prior information for the inference. Furthermore, a previous work shows that the inference may be facilitated in the presence of perturbations [63]. Along this line of work, our model showed that silencing the activity of a neural ensemble shifts the sequence to another ensemble, and the shift is controlled by the connectivity profile.

Previous studies have shown that sequences of activity might serve as a scaffold for local and downstream computations. For example, activity sequences in the mouse striatum enabled a read out of time in a downstream area [11], sequences in the mouse CA1 enabled the readout of cues and cue-specific elapsed time [7], and a recurrent network that encodes time through sequential activity can drive the representation of spatial information in a downstream network [65]. Since our findings demonstrate that neural sequences can generate a diversity of sequential dynamics in downstream regions, our results have the potential of unmasking the mechanisms underlying computations that are mediated by flexible and compositional sequences across circuits.

## Supporting information

Supplementary material with analytical derivations

## ACKNOWLEDGMENTS

We thank Sara Solla for valuable discussions and for her helpful feedback about the manuscript. We also thank Martina Acevedo for discussions about connectivity motifs. This work was supported by the Norwegian University of Science and Technology (SGC), by the Research Council of Norway through its Centres of Excellence scheme, project number 332640 (SGC), through a Researcher Project for Early Career Scientists, project number 102744100 (SGC), by the Kavli Foundation (SGC), and by two exchange grants awarded to SGC: IBRO Exchange Fellowship awarded and EMBO Scientific Exchange Grant.

## AUTHOR CONTRIBUTIONS

Visualizations: LMB; Simulations and analytical derivations: LMB, MKN, SNHR, SGC; Manuscript writing: LMB and SGC with input from all authors; Conceptualization and supervision of the project: CC and SGC; Funding acquisition: SGC.

## DECLARATION OF INTERESTS

The authors declare no competing interests.

## DECLARATION OF AI TECHNOLOGIES IN THE WRITING PROCESS

The authors used ChatGPT for improving the readability of the manuscript and for checking the grammar. The authors take full responsibility for the content of the article.

## RESOURCE AVAILABILITY

Requests for further information and resources should be directed to and will be fulfilled by the lead contact, Soledad Gonzalo Cogno.

### Materials availability

This study did not generate new unique reagents

### Data and code availability

All codes will be made publicly available upon publication. No new data were collected for this study.

## FIGURE TITLES AND LEGENDS

**Supplementary Figure 1:**
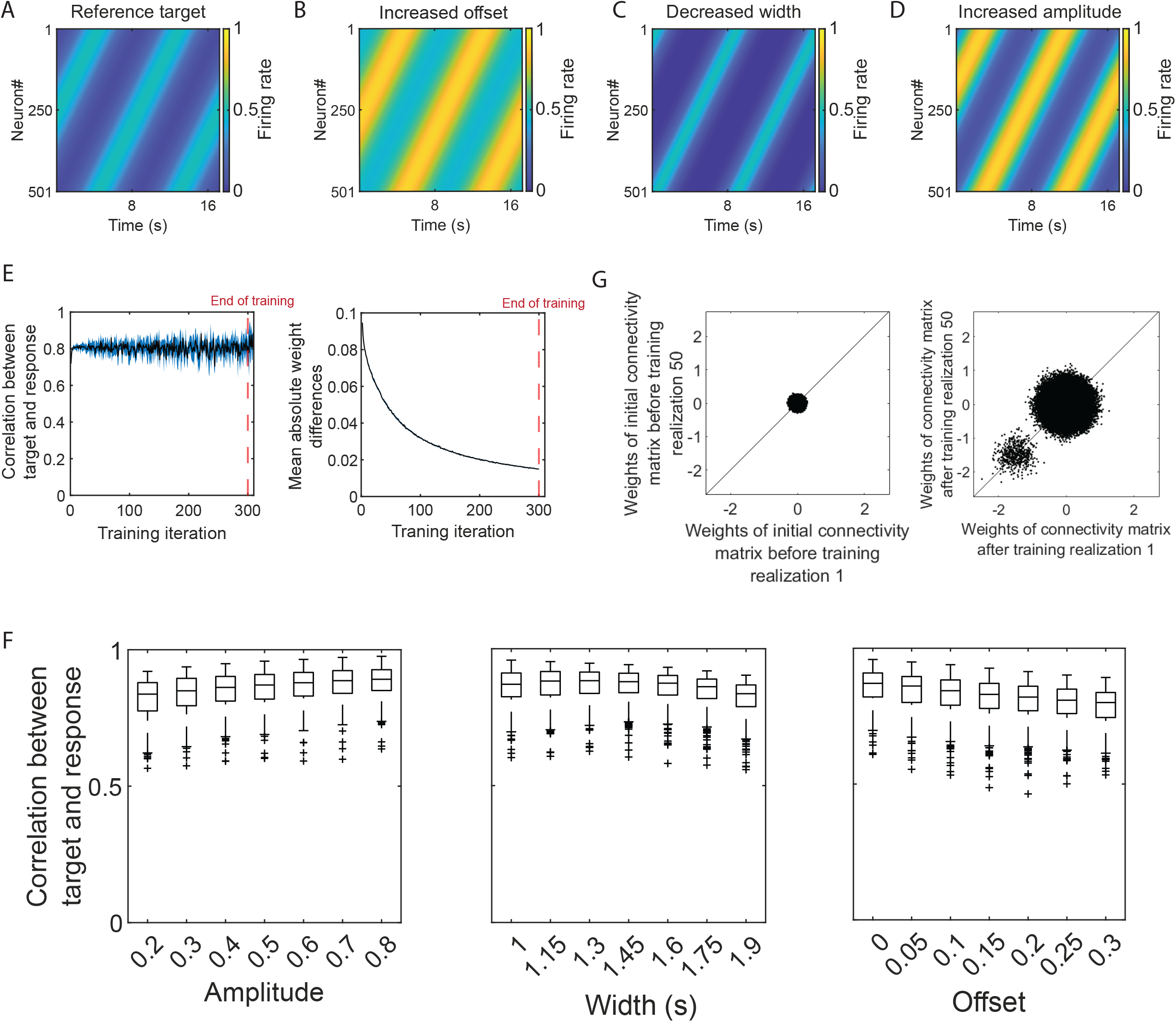
A recurrent neural network is successfully trained to generate sequences with different properties. **A**, Reference target sequences for an RNN of 501 units. Amplitude = 0.5, Width = 1.7 s, Offset = 0. Symbols and colours as in Figure 1B. **B-D**, Target sequences with the same parameters as in A except for Offset = 0.4 (B), Width = 1 s (C), Amplitude = 1 (D). **E**, Left: Mean correlation (black) between the target sequences and the RNN response calculated over 50 ‘training realisations’ and depicted as a function of the ‘training iterations’ (300 in total). The error bars (blue) indicate the standard deviation across training realisations. The end of training is marked with a red dashed line. We froze the weights from the last training iteration, that is training iteration 300, and let the network run for ten additional ‘test iterations’. We show the correlation values for the test iterations right next to the training iterations. Right: Mean absolute difference between weights of the connectivity matrix in successive training iterations. The mean was calculated over 50 training realisations. Symbols as in the left panel. Same sequence parameters as in panel A. **F**, Pearson correlation between the target sequences and the RNN response as a function of parameters of the target sequences. Each boxplot shows the correlation values across 50 training realisations. The horizontal line indicates the median, the bottom (top) extreme of the box indicates the 25^th^ (75^th^) percentile, the vertical lines indicate the maxima and minima, and the crosses outliers. Left: Width = 1 s, Offset = 0, 0.2 ≤ Amplitude ≤ 0.8. The training performance increases with increasing amplitude values. Middle: 1 s ≤ Width ≤ 1.9 s, Offset = 0, Amplitude = 0.5. The training performance remains approximately the same across width values. Right: Width = 1 s, 0 ≤ Offset ≤ 0.3, Amplitude = 0.5. The training performance decreases with increasing offset values. **G**, Left: Initialization of weights corresponding to training realisation 1 vs training realisation 50. These are the weights before training. Right: Same as left but after training. The values deviate from the diagonal (black line), indicating differences between the two connectivity matrices obtained after training.

**Supplementary Figure 2:**
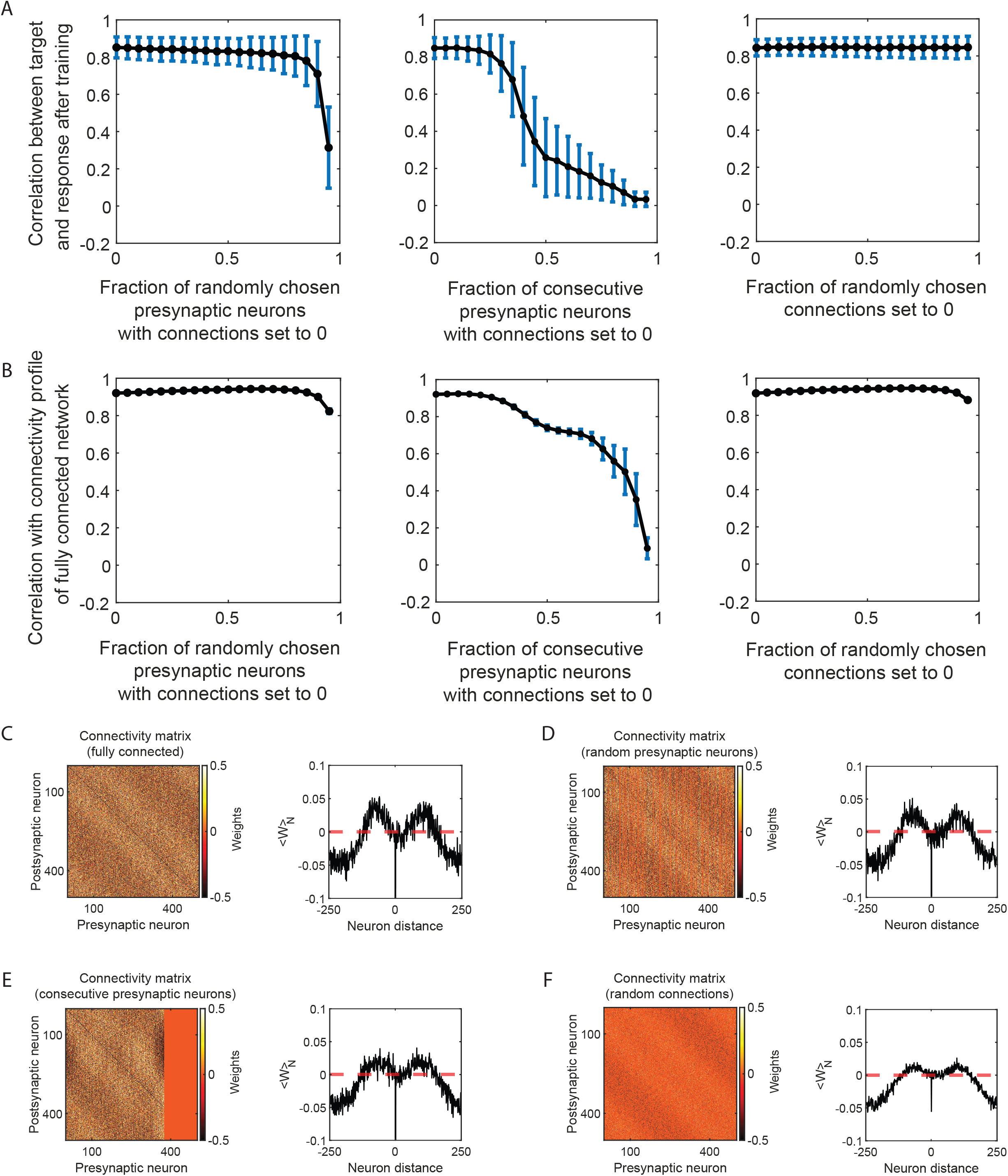
The similarity across connectivity profiles holds for sparse connectivity. **A**, Left: Mean correlation (black) between the target sequences and the RNN response calculated over 10 test iterations and depicted as a function of the fraction of presynaptic neurons from which the outgoing connections were set to 0 before training. The presynaptic neurons were randomly chosen, and those weights were not trained. The error bars (blue) indicate the standard deviation across test iterations grouped from 50 training realisations. Amplitude = 0.5, Width = 1.7 s, Offset = 0. The outgoing synapses of up to 95% of randomly selected neurons can be set to 0 and the network will still learn the target sequences. Middle: Same as the left panel, but now the presynaptic neurons are located consecutively in the sequence ordering. The outgoing weights of up to 35% of consecutive presynaptic neurons can be set to 0 and the network will still learn the target. Right: Similar to the left panel, but now randomly chosen synapses are set to 0. Unlike in the other two panels, now a single neuron can have outgoing synapses that are set to 0 and other outgoing synapses whose weights are trained. Up to 95% of the synapses can be set to zero and the network will still learn the target sequences. **B**, Similar to A, but for the correlation between the connectivity profile calculated on the mean connectivity matrix after training when the RNN is fully connected, and the connectivity profile computed on the connectivity matrices after 50 training realisations when the connectivity was sparse. The black line indicates the mean correlation across training realisations and the blue error bars the standard deviation. If the training was successful (see A), the correlation between the connectivity profiles of the sparse and the fully connected network was high. **C**, Left: Connectivity matrix corresponding to the target sequences in Figure 1B, Amplitude = 0.5, Width = 1.7 s, Offset = 0. The RNN was fully connected. Right: Connectivity profile (black) corresponding to the matrix on the left. Symbols and colours as in Figure 1C. **D**, Similar to C, but the outgoing connections of 25% of randomly chosen presynaptic neurons was set to 0. Correlation between the connectivity profile in C, and the connectivity profiles obtained from 50 training realisations when the network had this degree and type of sparseness: mean ± STD = 0.930 ± 0.004. **E**, Similar to C, but the outgoing connections of the last 25% of presynaptic neurons in the sequence ordering was set to 0. Correlation between the connectivity profile in C, and the connectivity profiles obtained from 50 training realisations when the network had this degree and type of sparseness: mean ± STD = 0.910 ± 0.005. **F**, Left: Similar to C, but 85% of all synapses were randomly chosen and set to 0. Correlation between the connectivity profile in C, and the connectivity profiles obtained from 50 training realisations when the network had this degree and type of sparseness: mean ± STD = 0.940 ± 0.003.

**Supplementary Figure 3:**
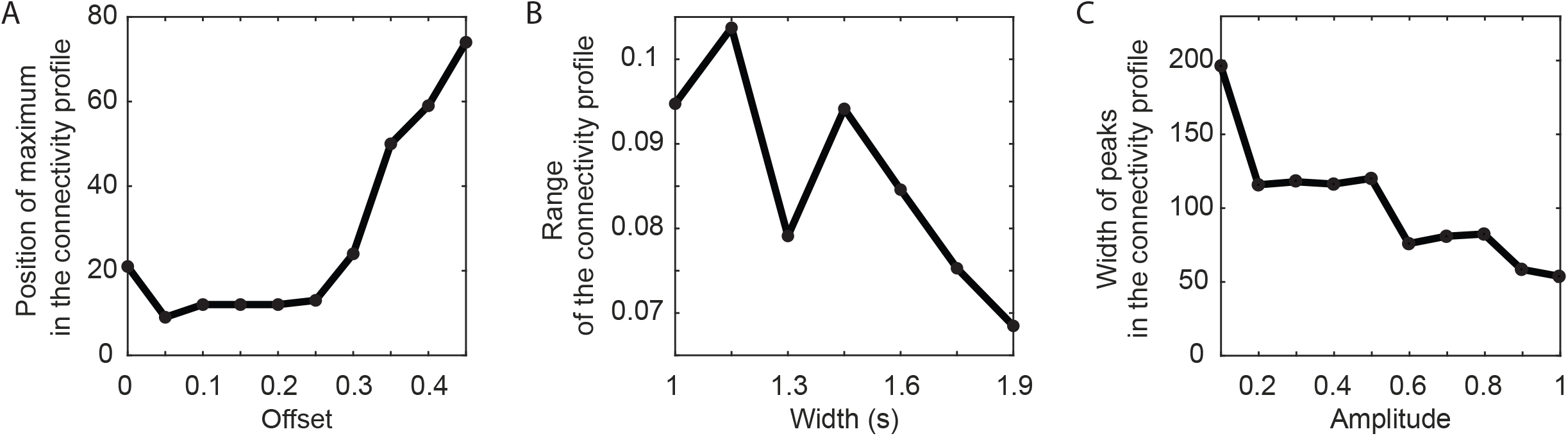
The connectivity profile changes smoothly with changes in the parameters of the target sequences. **A**, Position of the maximum in the connectivity profile, measured in distance between neurons, as a function of offset values. Amplitude = 0.5, Width = 1 s. We used the position of the maximum as a quantification of the degree of asymmetry of the connectivity profile. **B**, Range of the connectivity profile, defined as the difference between its maximum and minimum, as a function of width values. Amplitude = 0.5, Offset = 0. **C**, Mean width of the peaks in the connectivity profile as a function of amplitude values. Width = 1 s, Offset = 0. We calculated the width of the peaks as a proxy for the quantification of the frequency of oscillation in the connectivity profile, under the assumption that the wider the peaks, the smaller the frequency and vice versa.

**Supplementary Figure 4:**
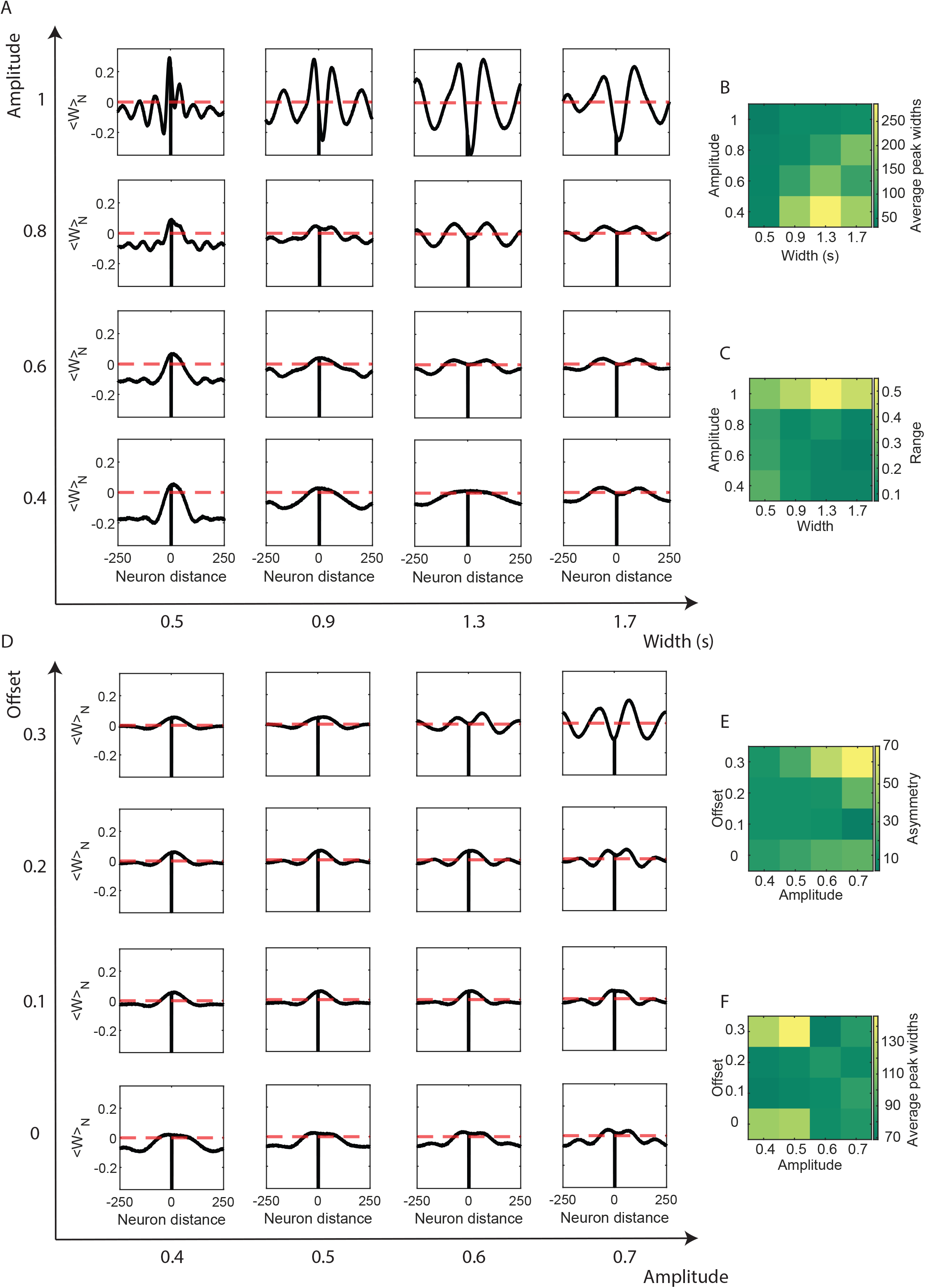
The connectivity profile smoothly maps changes in pairs of properties of the target sequences. **A**, Connectivity profiles corresponding to the connectivity matrices that generate sequences with varying amplitude and width values. Offset = 0. Symbols and colours as in Figure 1C. **B**, Heat map of the average peak width calculated on the connectivity profiles shown in A. Yellow indicates broad peaks, dark green indicates narrow peaks. The peaks are narrower, and therefore the oscillatory frequency higher, in connectivity profiles with high amplitude values. Offset = 0. **C**, Heatmap of the range of the connectivity profiles shown in A. The range is defined as the difference between the maximum and minimum. Yellow indicates high range, dark green indicates low range. The range decreases with increasing width values except when the amplitude is close to 1, when the range remains approximately the same. Offset = 0. **D**, Similar to A, but for different combinations of offset and amplitude values. Width = 1 s. **E**, Heatmap showing the asymmetry of the connectivity profiles in D, quantified through the position of the maximum. Yellow indicates high asymmetry, dark green indicates low asymmetry. The asymmetry increases in connectivity profiles with high offset and amplitude values. Width = 1 s. **F**, Similar to B, but for the connectivity profiles in D. The average peak width decreases with increasing amplitude values of the target sequences. Width = 1 s.

**Supplementary Figure 5:**
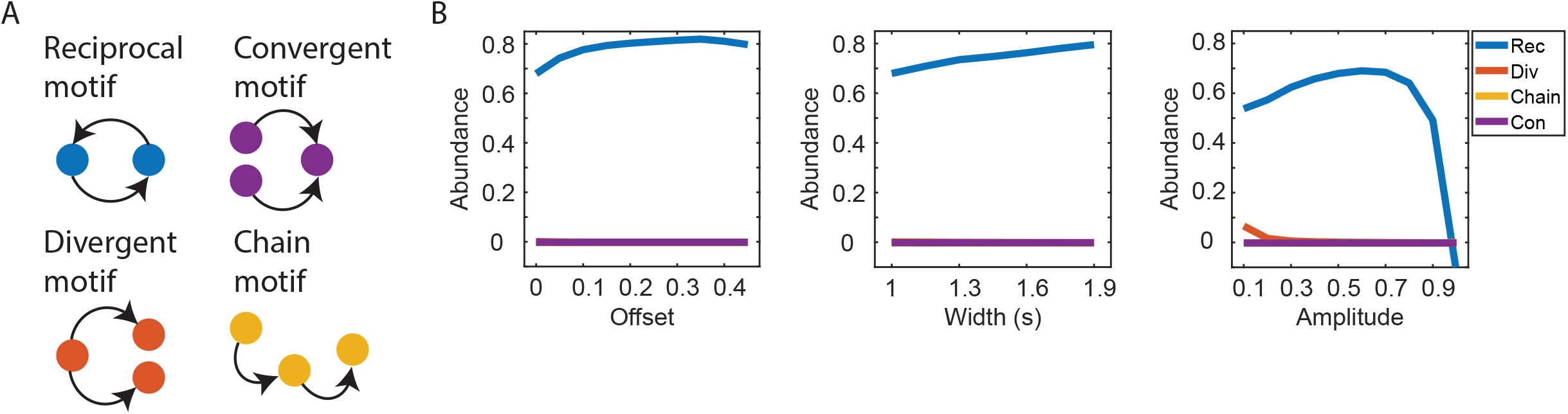
Second order statistics of the connectivity matrix map changes in the parameters of the target sequences. **A**, Schematics of connectivity motifs. We focused on: reciprocal motifs (top left), in which two neurons are mutually connected, divergent motifs (bottom left), in which one neuron connects to two other neurons, convergent motifs (top right), in which two neurons connect to the same neuron, and chain motifs (bottom right), in which one neuron is connected to a second neuron, which is connected to a third neuron **B**, Abundance of motifs as a function of the target sequences parameters: offset (left), width (middle), and amplitude (right). The abundance of the reciprocal motif is indicated in blue, the divergent motif in orange, the chain motif in yellow, and the convergent motif in purple. Left: The abundance of the reciprocal motif increases with increasing offset values. Amplitude = 0.5, Width = 1 s. Middle: The abundance of the reciprocal motif increases with increasing width values. Amplitude = 0.5, Offset = 0. Right: The abundance of the reciprocal motif increases with increasing amplitude values up to 0.7, after that value it decreases. Width = 1 s, Offset = 0. The chain, divergent and convergent motifs are not present in any of the connectivity matrices.

**Supplementary Figure 6:**
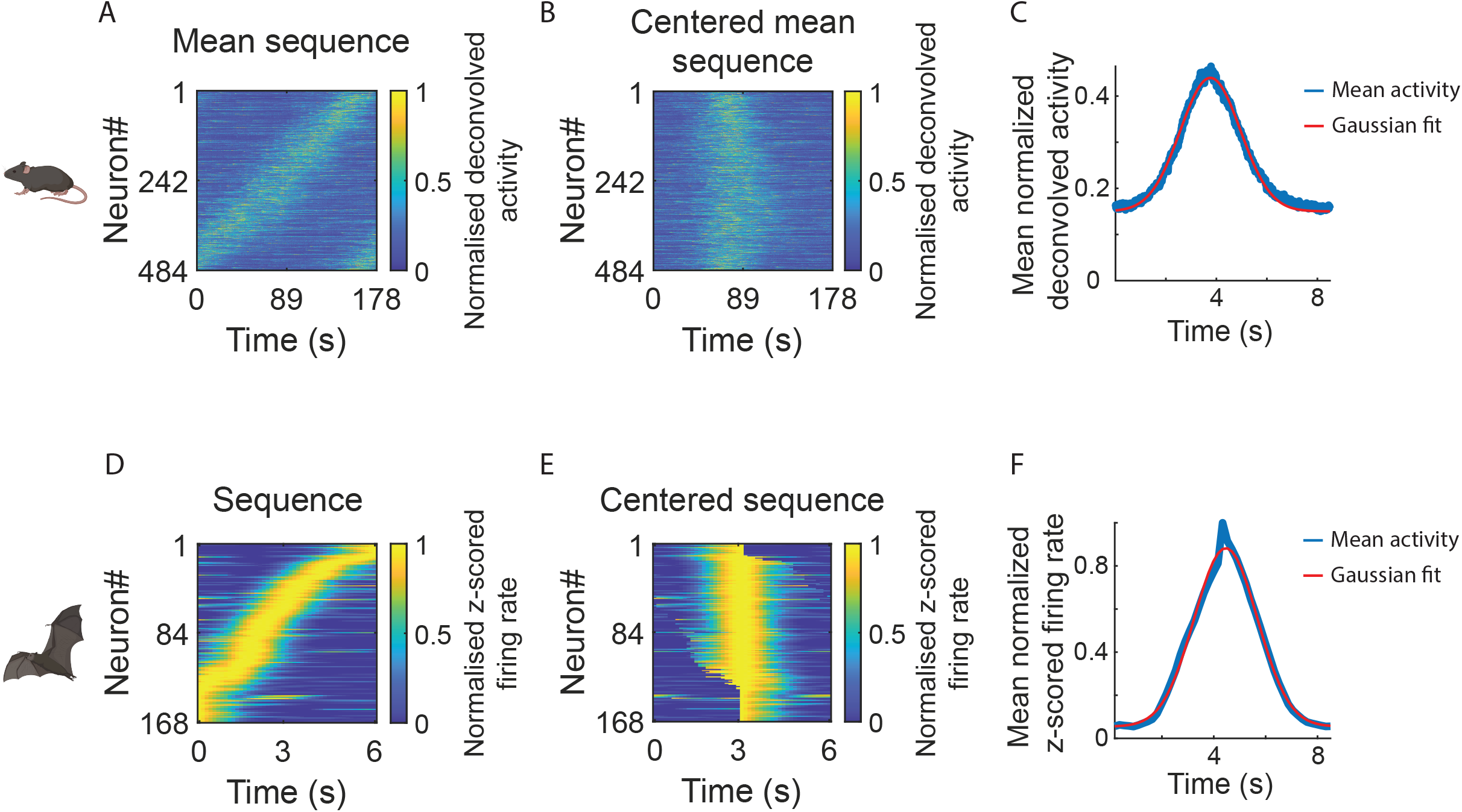
Procedure for fitting experimental data. **A**, Mean sequence of activity calculated over 24 periodic sequences available in experimental data session 7 from mouse #60584, published in Gonzalo Cogno et al., 2024. For computing the mean sequence, we first scaled the duration of each individual sequence so that they would all have the same duration: 177,7 s. We next calculated the mean activity of each neuron across sequences. Symbols and colours as in Figure 1B. **B**, We circularly shifted the activity of each neuron in panel A so that the activity of all neurons peaked at the same time bin. Symbols and colours as in Figure 1B. **C**, We first rescaled the centered mean sequence, depicted in panel B, so that the activity of each unit was 8.5 s long, which is the duration of the sequences in the simulated data. Then, we calculated the mean activity across neurons (blue). Finally, we fit a Gaussian (red) to the rescaled mean activity. Obtained fitted parameters: Amplitude = 0.29, Width = 1.68 s, Offset = 0.15. Goodness of fit (R^2^): 0.99. **D**, Sequential activity of time cells recorded from the bat CA1. Symbols and colours as in Figure 1B. **E**, Similar to B, we centered the sequence by circularly shifting each neuron’s activity from D. **F**, Same as C but for the centered sequence of time cells depicted in E. Obtained fitted parameters: Amplitude = 0.83, Width = 1.72 s, Offset = 0.05. Goodness of fit (R^2^): 0.99.

**Supplementary Figure 7:**
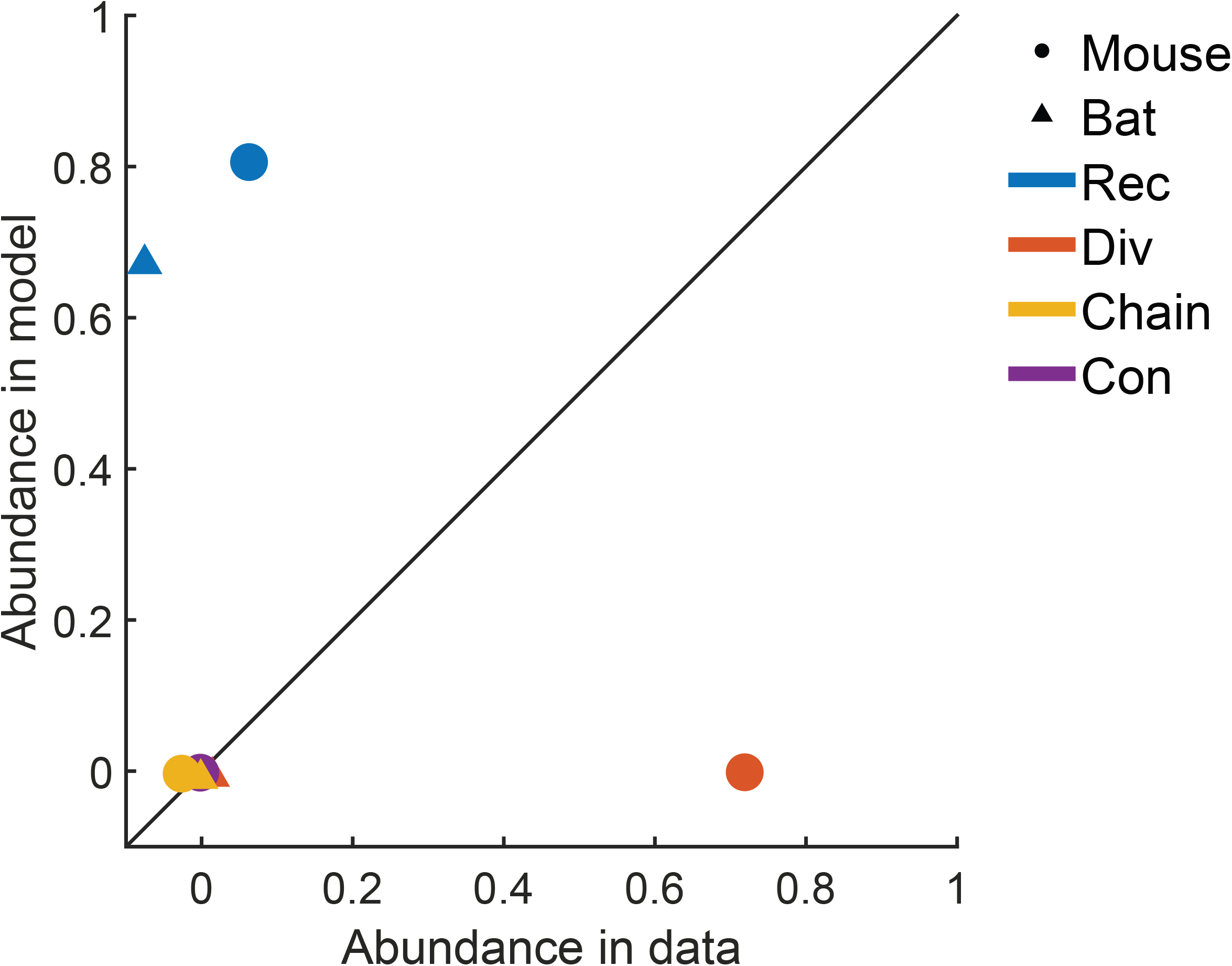
Comparison of connectivity motifs between experimental and synthetic data. Abundance of connectivity motifs obtained after training the RNNs on experimental data, as a function the same motifs obtained after training the RNNs on synthetic targets that best approximate the experimental data. The abundance of motifs corresponding to mouse (bat) data are indicated with full circles (triangles). Reciprocal motifs are indicated in blue, divergent motifs in orange, chain motifs in yellow, and convergent motifs in purple. The black line indicates the identity line. Chain and convergence motifs are absent in all connectivity matrices. There is a mismatch between motifs calculated on experimental data and on synthetic data.

**Supplementary Figure 8:**
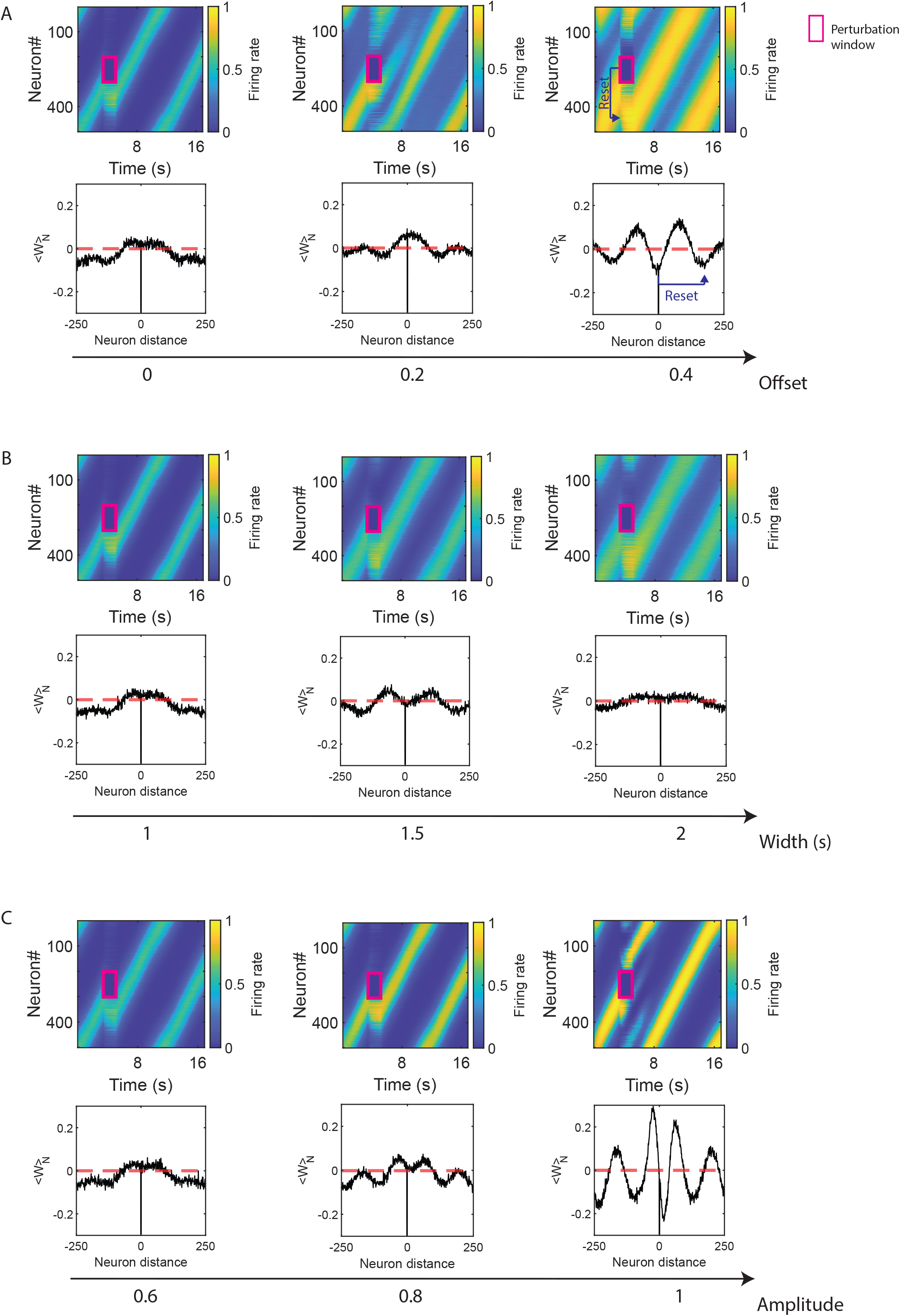
After silencing perturbations the sequences reset, and the ensemble to which they reset is controlled by the connectivity profile. **A**, Top: RNN response as a function of offset values. The perturbation time window, and the ensemble of neurons whose activity was set to zero, is indicated with a magenta rectangle. The perturbation induces a reset of the sequence for all offset values of the target sequences. Amplitude = 0.6, Width = 1 s. Bottom: Connectivity profiles calculated on the connectivity matrix that generates the unperturbed sequential dynamics corresponding to the top panels. The ensemble to which the sequence resets is as far away from the silenced ensemble as the position, measured in distance between neurons, of the minimum in the connectivity profile. In the right panel, the blue arrow indicates the reset in the response (top) and in the connectivity profile (bottom). Symbols and colours as in Figure 1B,C. **B**, Similar to A, but for increasing width values of the target sequences. Amplitude = 0.6, Offset = 0. **C**, Similar to A, but for increasing amplitude values of the target sequences. Width = 1 s, Offset = 0. For amplitude = 1 (right) parallel and transient sequences emerge due to the perturbation. This is due to the presence of oscillations, and therefore multiple minima, in the connectivity profile. Symbols and colours as in Figure 1B, C.

**Supplementary Figure 9:**
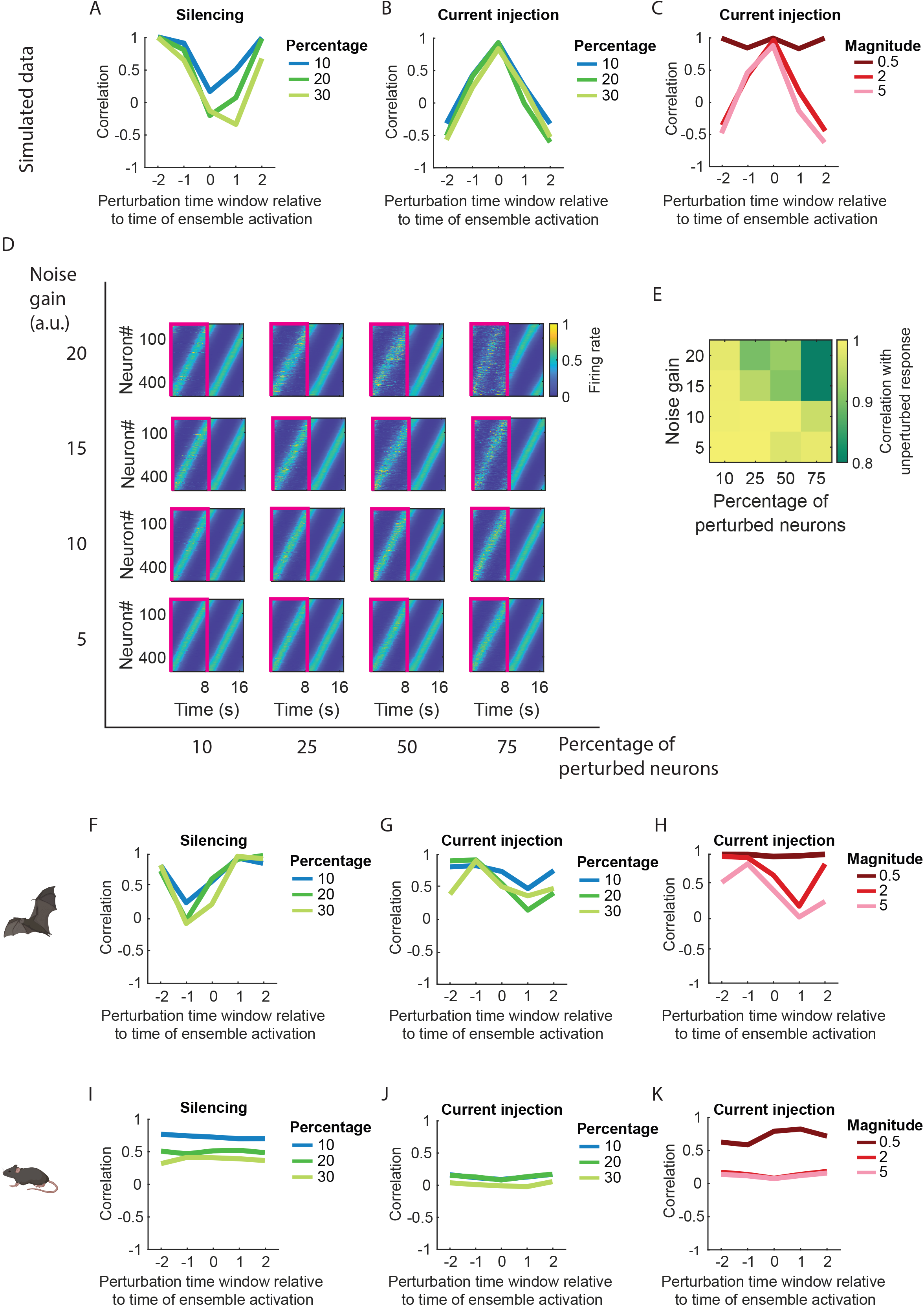
Effect of perturbations on the RNN dynamics. **A**, Correlation between the unperturbed network activity and the network activity after the perturbation was released. The correlation is depicted as a function of the time window in which the perturbation was applied, measured relative to the time in which the unperturbed sequential activity unfolded. If the perturbation was applied to an ensemble that was not engaging in the sequence, the perturbation time window relative to the time of ensemble activation was close to 2 or -2. If the perturbed ensemble was engaging in the sequence, the perturbation time window was 0. Ensembles of size corresponding to 10% (blue), 20% (dark green), or 30% (light green) of all neurons in the RNN (501 units) were silenced during a time window of 1.7 s. If the perturbation was applied to an ensemble that was not engaging in the sequence, the correlation was close to 1. If the perturbed ensemble was engaging in the sequence, the correlation dropped close to zero. Perturbing a higher fraction of neurons (shades of green) led to more negative correlations. Parameters of the target sequences: Amplitude = 0.6, Width = 1 s, Offset = 0. **B**, Similar to A, but for the current injection perturbation. Magnitude of the step current = 3. Injecting current into neurons that were not active in the sequence led to a negative correlation, corresponding to a reset of the sequence. Injecting current into neurons engaging in the sequence led to a correlation close to 1 corresponding to an unaffected progression of the original sequence. The effects are independent of the fraction of neurons that were perturbed. **C**, Similar to B, but each line indicates different magnitudes of the step current (0.5 in brown, 2 in red, 5 in pink). For very low magnitudes of injected current the correlation was close to 1, and hence the perturbation did not have any effect on the progression of the sequences. For higher magnitude values, the effect of the perturbation depended on the time window of current injection, similar to B. Ensemble size = 20% **D**, Different percentages of randomly selected neurons (indicated on the x axis, N = 501) were perturbed by injecting noise of different gains (indicated on the y axis). The time window of perturbation is 8.5 s long, equal to the duration of the first sequence, and is marked in magenta. The sequences always continue uninterruptedly after the perturbation was released. Amplitude = 0.5, Width = 1 s, Offset = 0. Symbols and colours as in Figure 1B. **E**, Combinations of noise gain (y axis) and percentage of perturbed neurons (x axis). The correlation between the unperturbed network response and the network response after the perturbation was released is indicated in shades of green. **F-H**, Same as A-C but for a network trained on bat experimental data. For a current injection of small magnitude the correlation value was always close to 1, and hence the sequential dynamics was not affected by the perturbation independently of the time window at which the current was injected. The correlation values decreased with increasing magnitudes of injected current. **I-K**, Same as A-C but for a network trained on mouse experimental data. The correlation was close to 0 due to an attractor state that the network settled into after the perturbation was released. For some training realisations, however, the network did not converge to the attractor state and instead the sequence unfolded uninterruptedly after the perturbation was released (not shown).

**Supplementary Figure 10:**
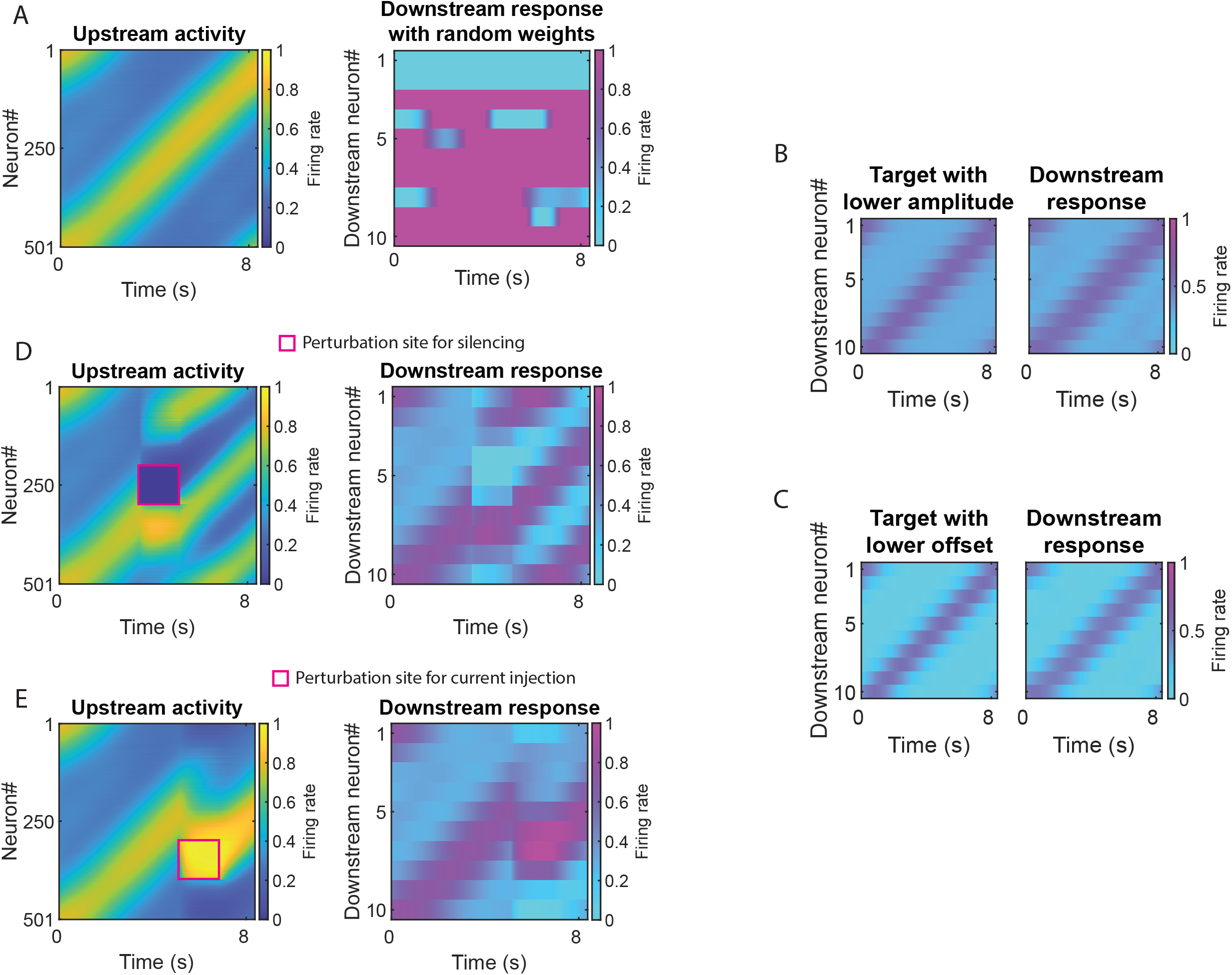
Downstream sequences are constrained by the properties of the upstream sequences. **A**, Left: Upstream sequential dynamics (501 neurons). Symbols and colours as in Figure 1B. Right: Response of the downstream network (10 neurons) when all upstream neurons are randomly connected to all downstream neurons. When the weights are random the downstream network does not generate sequential dynamics despite receiving sequential input from the upstream network. Colours as in Figure 5B. **B**, Left: Downstream target sequence. Parameters of downstream target sequence: Amplitude = 0.3, Width = 1 s, Offset = 0.3. Parameters of upstream sequence: Amplitude = 0.5, Width = 1 s, Offset = 0.3. The amplitude of the downstream target is lower than the amplitude of the upstream sequence (Figure 5B left). The upstream sequence and the downstream target are both 8.5 s long. Right: Response of downstream network after training using the target on the right panel. The target and the response are similar (Correlation: 0.99). Colours as in Figure 5B. **C**, Similar to B, but now the offset of the downstream target is lower than the offset of the upstream sequence. Parameters of downstream target: Amplitude = 0.5, Width = 1 s, Offset = 0.05. Parameters of the upstream sequence as in panel B. The upstream sequence and the downstream target are both 8.5 s long. The target and the response are similar (Correlation: 0.99). **D**, Left: Upstream sequential activity (501 neurons). We perturbed the upstream sequence by silencing the activity of 100 neurons, in a time window of duration = 1.7 s, indicated in magenta. Symbols and colours as in Figure 1B. Right: Response of the downstream network receiving as input the activity on the left. The weights between the upstream and the downstream network were trained using as input the unperturbed response shown in panel A, left. The downstream neurons showed a similar reset in their activity as observed in the upstream neurons. Parameters of upstream sequence: Amplitude = 0.6, Width = 1 s, Offset = 0.3. The upstream sequence and the downstream target are both 8.5 s long. Colours as in Figure 1B, 5B. **E**, Same as D but instead of silencing, additional current was injected into the ensemble of neurons, and at the time window, indicated in magenta. Magnitude of step current: 3. The downstream neurons showed a similar reset in their activity as the upstream neurons.

## METHODS

### Recurrent neural network model

To characterize the set of connectivity matrices that generate sequential dynamics, we constructed rate-based recurrent neural network (RNN) models, and we trained the weights in the RNNs so that the entire network would generate sequential activity. Our approaches for designing and training the RNNs are similar to those in Rajan et al., 2016.

We worked with RNNs that were trained on simulated sequences of activity, or on neural sequences that were experimentally recorded. When we considered simulated sequences the network consisted of *N* = 501 units, as we show in figure 1, 2, 4, and 5. When we considered sequences of activity that were experimentally recorded, as we show in figure 3, the network consisted of *N* = 170 neurons to match the experimental data recorded from the bat hippocampal CA1[13], or of *N* = 484 neurons to match the experimental data recorded from the mouse medial entorhinal cortex [12] (MEC).

Each neuron is represented by an activation variable *h*_*i*_, *i* = 1, . ., *N*. To obtain the firing rate *r*_*i*_ for each neuron *i*, we passed the activation variable through a sigmoid activation function: 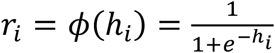. Because we used a sigmoid as the non-linearity 0 ≤ *r*_*i*_ ≤ 1 ∀*i*, i.e., the rates were bounded between 0 and 1.

The network connectivity is represented by a connectivity matrix *W* ∈ ℝ^*N*×*N*^. Each entry *w*_*ij*_ represents the weight from presynaptic neuron *j* to postsynaptic neuron *i* with *i, j* = 1, . ., *N*.

The temporal progression of the activation variable *h*_*i*_ of each neuron is determined by

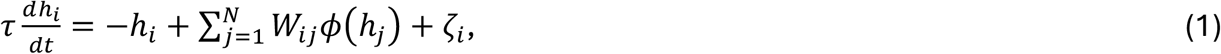

with *τ* = 0.3224*s*.

The noise *ζ*_*i*_ was generated as

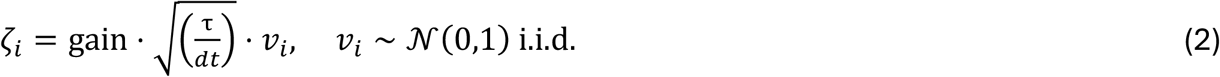

with gain = 0.5 and time step *dt* = 0.0081s. The results of the paper remain unchanged for *dt* = 0.1 s and *dt* = 0.2 s (not shown).

Let *R*(*t*) = (*r*_1_(*t*), …, *r*_*N*_(*t*))^*T*^ be the vector of firing rates at time *t*, and *H*(*t*) = (*h*_1_(*t*), …, *h*_*N*_(*t*))^*T*^ the vector of activation variables at time *t*, both of size *N*. To integrate the network activity, we used the Euler method with an integration step *dt* = 0.0081*s* as follows

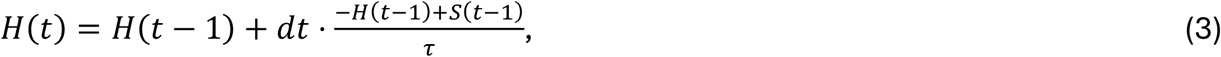

with *S*(*t*) = *WR*(*t*) + *ζ*(*t*)

and *ζ*(*t*) defined as the vector *ζ*(*t*) = (*ζ*_1_(*t*), …, *ζ*_*N*_(*t*)), of size *N*.

We explain how we initialized and how we trained the components of *W* below, in the section ‘FORCE-training of the RNNs’.

### Design of simulated target sequences

To train the RNNs we first constructed a target activity for each neuron such that, after training, each neuron would generate activity resembling its target. The target firing rate of each individual neuron *i* = 1, …, *N*, is given by

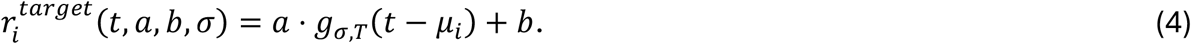

Let *T* be the duration of the neural sequence, with *T* = 8.5 *s* in the case of synthetic sequences that we describe here, then 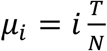, and *g* _*σ,T*_ is a wrapped Gaussian given by:

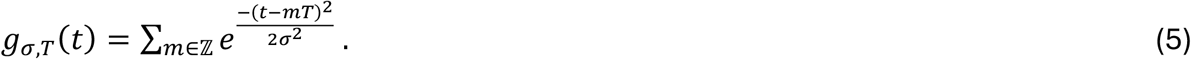

We simulated 18 periodic sequences by evaluating the target activity over a time interval of length 18*T* (note that the Gaussian repeats over time) in a population of *N* = 501 neurons. The period of the sequences was *T*. The ordering of the cells in the sequence is given by *μ*_*i*_, i.e., based on the time at which the firing rate of the cells peaks. We used a bin size = 0.129 s. This choice of bin size is so that the synthetic data is generated at a similar temporal resolution to the one we have in the experimental data (see ‘Experimental target sequences’).

We normalized the activity of each unit by 1 Hz, therefore the offset and the amplitude are dimensionless. The width has units of time.

We defined the ‘target sequences’ as the set of all target firing rates: 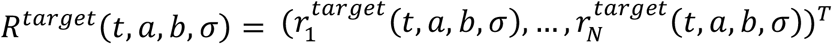, where *R*^*target*^(*t, a, b, σ*) is a vector of size *N*, that also depends on the parameters *a, b, σ*. The combination of ‘amplitude’ *a*, ‘width’ *σ*, and ‘offset’ *b* determines the properties of the sequence. The amplitude took values between 0.2 and 1, the width between 1 and 1.9, and offset between 0 and 0.45. For example, the sequences in Figure 1B correspond to the parameters *a* = 0.5, *σ*= 1.7, and *b*= 0. The target sequences were 153 s long, for a total 18 periodic sequences of *T* = 8.5 *s*.

We calculated the ‘target activation variables’ of each neuron, denoted as 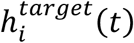, by applying the inverse of the sigmoid activation function to the target firing rates of each neuron:

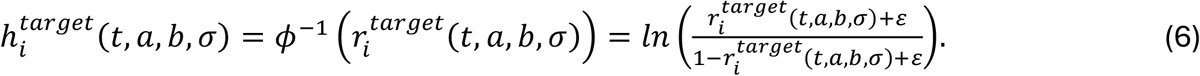

with *ε*=0.01, to avoid dividing by 0 if 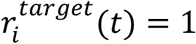 at any time point 0 ≤ *t* ≤ 18 ⋅ *T*. Similarly, as we did for the target sequences, we introduced the ‘network target current’ as 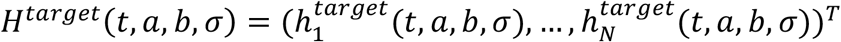.

As mentioned above, the model integration time step is *dt* = 0.0081 s and the target sequences are constructed using a *bin size* = 0.129 s. The results of the paper remain unchanged for other choices and combinations of *dt* and *bin size*: with *dt* ∈ {0.0081 s, 0.1 s, 0.2 s} and *bin size* ∈ {0.1 s, 0.129 s, 0.2 s} (not shown).

The mismatch between *dt* = 0.0081 s and *bin size* = 0.129 s implies that for each time bin of the target sequences, there are approximately 16 time steps of the dynamics of the model. This is relevant for the training procedure (‘FORCE-training of the RNNs’). The network activity was initialized with *R*^*target*^(*t* = 0) and *H*^*target*^(*t* = 0).

### Experimental target sequences

*Bat data*: We analysed publicly available electrophysiological recordings from bat hippocampal CA1, published in Omer, Las, and Ulanovsky 2023. In those experiments, two bats, the observer and the demonstrator, were in the same room. We only analysed neural data from the observer bat. The room was equipped with 3 landing balls, the start ball, ball A, and ball B. We analysed data from conditions in which the demonstrator bat was hanging motionlessly on either of the landing balls, and the observer bat landed on ball A or B until it took off. During the ‘hanging time intervals’, i.e., between landing and take off, hippocampal time cells fired in a sequential manner. The analysed data were z-scored firing rates of time cells averaged across hanging time intervals. Because the hanging time intervals varied in duration, the number of hanging time intervals that contribute to the calculation of the mean neural activity decreases with time after landing. To maximize the number of analysed cells and the number of hanging time intervals, we used only the first 6 seconds of the data that followed the observer bat’s landing on ball A or B. Consequently, the analysed dataset included the mean z-scored firing rates of 168 time cells over 6s after landings. To sort the time cells into a sequential order, we first smoothed their activity over 15 bins (bin size = 0.1*s*) using the MATLAB function “movmean” and afterwards, for each cell we identified the time point at which its firing rate peaked. We next sorted the neurons according to those time points, in an ascending order. Next, to compare the bat data to the experimental dataset recorded in mouse (see below), we downsampled the unsmoothed z-scored firing rate to a bin size of 0.129s and set all negative values to 0. Afterwards, we smoothed the neural activity over 15 time bins using the MATLAB function “movmean” and for each neuron divided the resulting signal by its maximum, leading to normalized activity with maximum value 1. Results remain unchanged for shorter and longer smoothing windows.

We used these data as targets for training an RNN with a matching number of 168 units, a time constant *τ* = 0.3224*s* and integration step *dt* = 0.0081*s* (same parameters as for the RNN trained on simulated data). Each unit received as target the activity of one cell.

Similar to what we did for the synthetic sequences, from the *R*^*target*^ computed as we describe above, in this case of size 168 × 1, we calculated *H*^*target*^. The network activity was initialized with *R*^*target*^(*t* = 0) and *H*^*target*^(*t* = 0).

#### Mouse data

We analysed experimental session 7 from mouse #60584 from the dataset reported in Gonzalo Cogno et al 2024. In that publicly available dataset, mice were head-fixed, placed on a running wheel in darkness conditions and did not receive any rewards. Calcium imaging recordings from the medial entorhinal cortex showed that the neural activity organized into ultraslow (frequency < 0.1 Hz) periodic sequences of activity. This session had 24 ultraslow periodic sequences. The deconvolved activity had a *bin size* = 0.129*s*. First, we smoothed the data over time bins with a moving mean and window of 120 bins using the MATLAB function “smoothdata”. To decrease runtime, we downsampled the data to obtain a bin size of 1.29 s. This does not change our results because the sequential dynamics are on a timescale of minutes. Afterwards, we normalized the deconvolved activity by dividing each neuron’s activity by its maximum value leading to normalized activity with maximum value 1.

Due to the periodicity of the sequences, for training the RNN we took 50 random snippets of 250 time bins (*bin size* = 1.29*s*; duration of each snippet = 322.5s) from the experimental data and used these snippets as targets to train 50 fully connected RNNs with initial random connectivity (see section ‘FORCE-training of the RNNs’). The networks consisted of 484 neurons, which is equal to the number of neurons in the experimental data. The time constant was *τ* = 0.3224*s* and the integration step *dt* = 0.0081*s* (same parameters as for the RNN trained on simulated data).

Similar to what we did for the synthetic sequences, from the *R*^*target*^ computed as we describe above, in this case of size 484 × 1, we calculated *H*^*target*^. The network activity was initialized with *R*^*target*^(*t* = 0) and *H*^*target*^(*t* = 0).

### Sequence ordering in synthetic and experimental target sequences

The sequence ordering, or the way neurons were labelled in the network, reflected the ordering according to which neural firing rate peaked as a function of time. In the synthetic data, the sequence ordering was obtained by sorting the time points at which the activity of the neurons peaked in an ascending order. In the experimental data, the ordering was obtained from the original datasets. In the CA1 bat dataset, similarly to what we did for synthetic sequences, the ordering was obtained by sorting the times at which the activity of each neuron peaked in an ascending manner. In the MEC mouse dataset, the sorting was obtained from applying PCA to the neural activity, and by sorting neurons based on PC1 and PC2 (see Gonzalo Cogno et al., 2024).

### FORCE-training of the RNNs

We trained the recurrent weights of the RNN using the targets described above. For each target *R*^*target*^, we performed 50 training realisations. In each realisation we randomly initialized the connectivity matrix *W*. Each weight *w*_*ij*_ was independently sampled from a standard Gaussian distribution and scaled by the factor 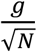, such that 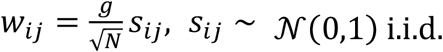, with *g*=1.5 for the simulated sequences and *g*=0.05 for the experimental sequences [33] and *N* as described above.

In the case of simulated sequences, and mouse experimental data, we used snippets of the target sequences as targets to train the network. To reduce computational time while preserving periodicity, each snippet encompassed two sequences. The snippet had a duration 2*T* (2*T* = 17*s* with bin size = 0.129*s* for simulated sequences, 2*T* = 322.5*s* with bin size *dt* = 1.29*s* for mouse experimental sequences). In each of the 50 training realisations, the initial time bin of the snippet was randomly chosen in the range [0,119.2] s for simulated sequences and [0, 2945.1] s for mouse experimental sequences. Note that for the simulated sequences, because those were perfectly periodic in time, all 50 targets were equivalent up to a shift in their sequence phase. In the case of bat experimental sequences, as in Figure 3C, the target consisted only of 1 sequence, i.e., had duration *T* (*T* = 6*s* for bat experimental sequence with bin size dt = 0.129s). Hence, the target was the same for all 50 training realisations.

In the following we refer to the snippet of the target sequences (simulated sequences, or experimental sequences) simply as ‘target’.

To train the RNNs we applied a variation of the FORCE training algorithm [34] that was introduced in Rajan et al. 2016 and which introduces a time dependent effective learning rate 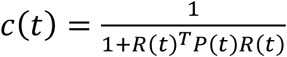. Each training iteration is as long as the target, i.e., 2086 time bins (= 16.9s) for simulated, 732 time bins (= 5.9s) for bat experimental sequences, and 39655 time bins (= 321.2s) for mouse experimental sequences. The integration time step in the model was always *dt* = 0.0081*s* leading to the set *T* = {*t* = *k* ⋅ *dt* ∣ 0 ≤ *k* ≤ *K*},*K* ∈ ℤ_≥0_, with *K* ≤ 2086 for simulated sequences, *K* ≤ 732 for bat experimental sequences, and *K* ≤ 39655 for mouse experimental sequences. We used 300 learning iterations during which the weights were adjusted according to equations 7-9 below, and 10 test iterations during which the weight matrix was held fixed.

As mentioned above, the target sequences are constructed using a bin size = 0.129*s* (synthetic sequences and bat experimental data) or bin size = 1.29*s* (mouse experimental data), whereas the model is integrated with a time step *dt* = 0.0081*s*. Consequently, each target time bin corresponds approximately to 16 integration time steps (simulated sequences and bat experimental sequences) or 160 integration time steps (mouse experimental sequences) in the model. Hence, every time bin of the target sequences (bin size = 0.129s) was used to train every 16^th^ integration time step in the model (dt = 0.0081s).

Let *T* = {*t* = *k* ⋅ *dt* ∣ 0 ≤ *k* ≤ *K*} with *K* as above be the set of all model integration time steps. The target values are available only at the subset *Z*_*m*_ ⊂ *T*

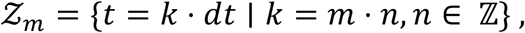

with fixed *m* = 16 for the simulated sequences and bat experimental sequences, *m* = 160 for the mouse experimental sequences and *n* ∈ ℤ. Denote the elements of *Z*_*m*_ as *z*.

The error is evaluated only at these time points *z* ∈ *Z*_*m*_:

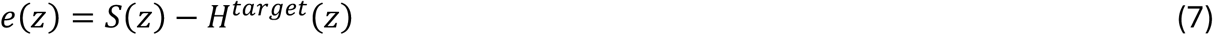

with *S*(*z*) and the noise term *ζ*(*z*) as introduced in section ‘Recurrent neural network model’.

The entries in the connectivity matrix *W* were updated according to a variation of FORCE learning [33, 34] as follows

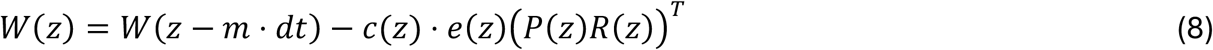

with

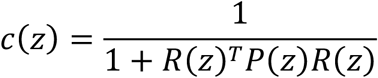

and

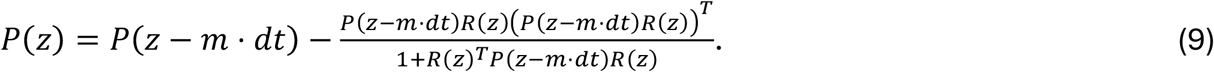

We used the correlation matrix *P* ∈ ℝ^*N*×*N*^which we initialized as *P*(0) = *p* ⋅ *I*, with *p*=0.08 a regularization term and *I* the identity matrix of size *N* × *N*.

For the time points *t* ∈ *T* ∖ *Z*_*m*_ as well as for every time point during the test iterations, the dynamics of the network were simulated without adjusting the connectivity matrix following equations 1-3.

Each of the 50 training realisations consisted of 300 training iterations, and in each training iteration the weights were updated at every time point *z* ∈ *Z*_*m*_. The initial weight matrix for iteration *q* was set equal to the final weight matrix from iteration *q* − 1, for *q* = 2, …, 300.

To evaluate the training performance within training realisations, we calculated the Pearson correlation between the dynamics of the RNN *R*(*z*) and the target. Additionally, we verified that the change of weights converged towards zero by calculating the mean absolute change of weights for each training iteration *q* = 1, …, 300, as follows

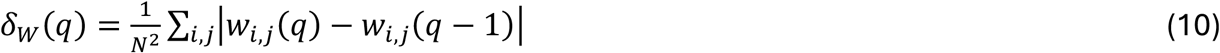

with *w*_*i,j*_(*q*) being the entry *i, j* with *i, j* = 1, …, *N* in weight matrix *W* after training iteration *q*.

### Quantification of the connectivity profile

After 50 training realisations, we obtained 50 independently trained connectivity matrices. Next, we computed the average connectivity matrix across the 50 resulting matrices. We tested that the average connectivity matrix generated sequences by examining its dynamics and confirming that it was sequential (Figure 1 F right). In the following, we refer to the average connectivity matrix simply as ‘mean connectivity matrix’. To quantify the structure of the connectivity matrices, we calculated their connectivity profiles as illustrated schematically in Figure 1C, denoted as *cp*_*W*_(*d*) with *d* being the distance between two neurons in the sequence ordering.

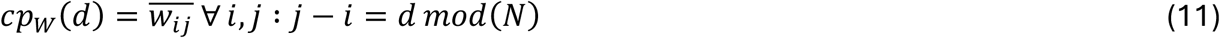

with *w*_*ij*_ being the weight between neuron *j* and *i*, and *N* the number of neurons in the network. We denote by *cp*_*W*_ the vector of the connectivity profile, i.e., 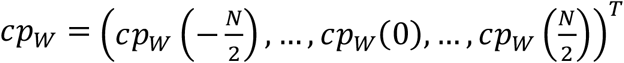.

To quantify how features of the connectivity profile were linked to properties of the sequential dynamics, we first preprocessed the connectivity profile by linearly interpolating the value of the self-connections based on the values of the connectivity profile at distance 1 and -1. To mitigate the effects of local minima, we smoothed the connectivity profile with a gaussian weighted moving average filter with a window size of 50 bins. To do this, we used the MATLAB function “smoothdata” with the method “gaussian” and window “50”. Whenever calculating extreme points, we used the MATLAB function “findpeaks” with minimum prominence 0.001.

- *Quantification of the oscillations in the connectivity profile:* We quantified the oscillations in the connectivity profile by calculating the width of the peaks, under the assumption that the more oscillations in the profile, the narrower the peaks would be. First, we identified all maxima in the connectivity profile. At each maximum, we set a reference line at height equal to half of their prominence and calculated their width at this reference line. Finally, we computed the mean width across all maxima.
- *Quantification of the range of the connectivity profile*: We defined the range of the connectivity profile as its maximum minus its minimum value:

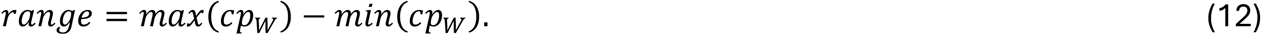
- *Quantification of the degree of asymmetry of the connectivity profile:* We defined the degree of asymmetry as the smallest absolute distance in the connectivity profile at which a local maximum occurs:

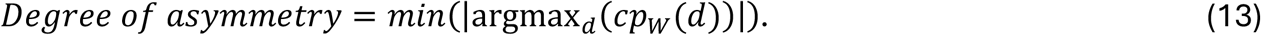

### Quantification of connectivity motifs

We calculated second order statistics of the connectivity matrix by computing the connectivity motifs [41]. First, we centered the connectivity matrix *W*

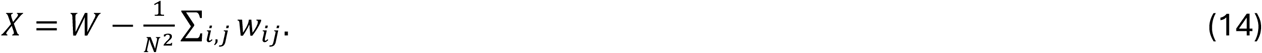

Next, we calculated the abundance of the following motifs: (i) the reciprocal motif, denoted as τ_rec_, where two neurons connect to each other, (ii) the divergent motif, denoted as τ_div_, where one neuron projects onto two different neurons, (iii) the convergent motif, denoted as τ_con_, where two neurons project onto the same third neuron, (iv) the chain motif, denoted as τ_chn_, where one neuron projects onto another neuron, which itself projects onto a third neuron (Supp. 5A).

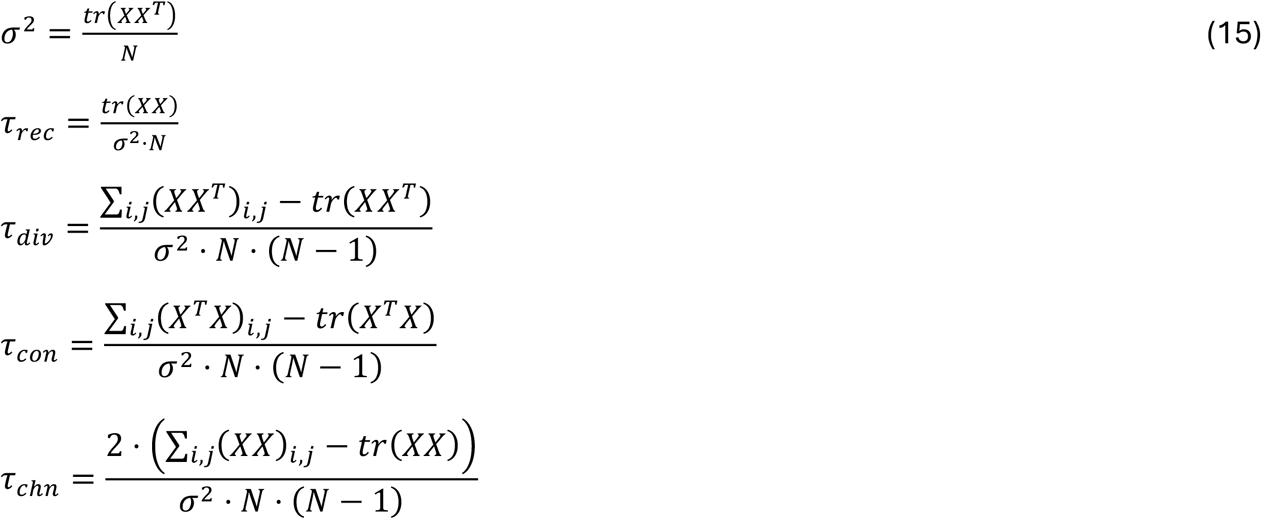

With *tr*(*X*) denoting the trace of matrix *X*.

### Incorporation of sparseness into the connectivity matrix

To investigate the robustness of the obtained connectivity profiles, we introduced sparseness into the connectivity matrix by setting up to 95% of all weights to 0.

#### a. Setting the outgoing weights of randomly selected neurons to 0

First, we set the outgoing connections of a random set of neurons, which included between 0 and 95% of all neurons, to 0, and we did not train these connections. Let *L* be the set of neurons whose outgoing connections are not set to 0, i.e., whose outgoing connections undergo training. We now define *R*_*L*_(*z*) as the rates of neurons in *L* at time *z*, i.e., *R*_*L*_ (*z*) is a vector of size *N*_*L*_, where *N*_L_ is the cardinality of *L* and *W*_*L*_ (*z*) is a matrix of size *N* × *N*_*L*_, which contains the outgoing weights of neurons in *L* at time *z* within a trainin*g* iteration, and *P*_*L*_ is the *N*_*L*_ × *N*_*L*_ correlation matrix between neurons in set L.

To train the RNN, the same procedure as described in equations 7-9 is followed. After calculating the error *e*(*z*) = *S*(*z*) − *H*^*target*^(*z*), the weights are adjusted as follows.

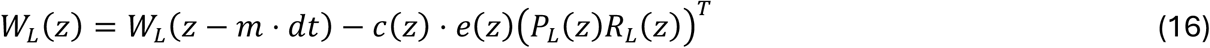

with

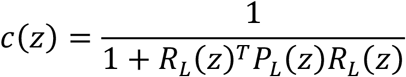

and

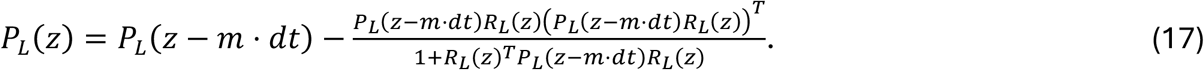

#### b. Setting the outgoing weights of neighbouring neurons in the sequence ordering to 0

In a different set of 50 training realisations, we proceeded as in the previous subsection ‘Setting the outgoing weights of randomly selected neurons to 0’, but now the cells in the set *L* were adjacent in the sequence ordering, instead of chosen in a random manner.

#### c. Setting randomly selected synapses to 0

Instead of setting all the outgoing weights from specific neurons to 0, we next set randomly chosen synapses to 0. Throughout our analyses we set between 0 and 95% of all synapses to 0 and did not train those weights.

The training procedure was similar to the learning rule as in the fully connected network (equations 7-9). Let *J*_*i*_ be the set of presynaptic neurons whose weights to neuron *i* = 1, …, *N*, undergo training, i.e., are not set to 0. In other words, *w*_*ij*_ ≠ 0 ∀*j ϵJ*_*i*_. Let 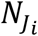 be the cardinality of *J*_*i*_. The weights between the neurons in *J*_*i*_ to neuron *i* are denoted as 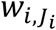, which is a vector of size 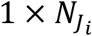. The rates of the neurons in *J*_*i*_ are denoted as 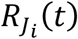, a vector of size 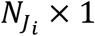. The weights in the RNN were updated according to the following equations.

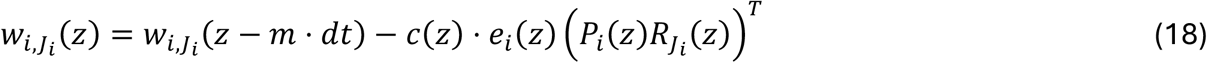

with

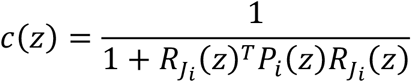

and

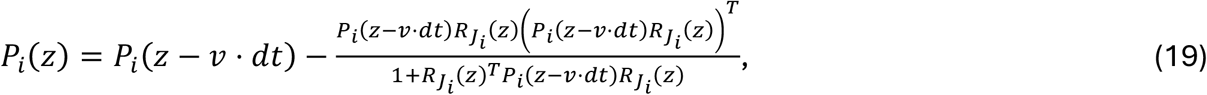

with *P*_*i*_, *i* = 1, …, *N*, now being a matrix of 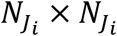 and 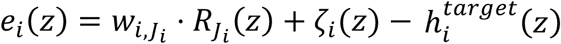 a scalar.

Note that the connectivity matrix *W* is initialized only once but the updates are applied per row *i* in columns *J*_*i*_ at each time point *z* and for each training iteration, and note that *P*_*i*_ is initialized independently for each neuron *i* (see ‘FORCE-training of the RNNs’).

### Generation of synthetic data that approximates experimental data

To generate simulated targets that match the experimental data as closely as possible, we first calculated for each dataset a ‘centered sequence’ (Supp. 6 B, E). We did it by taking the sequential activity and circularly shifting the activity of each individual neuron such that the sequence would now be centered, with the activity of all neurons peaking at the same time point. Next, we calculated the mean activity across neurons, which resulted in a signal that resembled a bump of activity, or ‘activity bump’, and fitted a Gaussian to it. Using the fitted parameters we generated synthetic sequences.

#### Calculation of activity bump in bat data

After preprocessing the data as in the section ‘Experimental target sequences’, we obtained the z-scored firing rates from 168 neurons over 6 s with bin size = 0.129 s (Supp. 6 D). We next circularly shifted each neuron’s activity, such that the activity of all neurons peaked at the same time bin (time bin 24) (Supp. 6 E). We refer to this as the centered sequence. Next, we rescaled the duration of the centered sequence (see subsection below), computed the mean across neurons, and obtained the activity bump in the bat data.

#### Calculation of activity bump in mouse data

We used the deconvolved activity from experimental data session 7 from mouse 60584 published in [12]. We identified all 24 sequences that were present that session. For further calculations, we excluded the first 4 sequences as those were interleaved with temporal epochs without sequential activity. From the remaining 20 sequences, we identified the one with the longest duration, which was 177.8 s, equivalent to 1378 time bins (bin size = 0.129 s). Next, we rescaled all other 19 sequences to this longest duration by linearly interpolating them so that each of them would also be 177.8 s long, and therefore would also encompass 1378 bins. Then, we calculated the mean across those 20 sequences (Supp. 6 A). We normalized the resulting sequence by dividing each neuron’s mean deconvolved activity by its maximum value. To denoise the neural activity, we defined a threshold for every neuron as the mean of its normalized deconvolved activity plus one standard deviation and set the activity of every time bin below the threshold to 0. Next, we calculated the centered sequence by circularly shifting the activity of each neuron so that the deconvolved activity of each neuron was centered at bin 689 (Supp. 6 B). Finally, after rescaling the duration for the sequence (see next subsection) we computed the mean across neurons to obtain the activity bump in the rodent data.

#### Procedure for fitting a Gaussian to the bump of activity calculated on experimental data

Because all synthetically generated sequences were 8.5 s long, we first rescaled the duration of the centered sequence to 8.5 s. Afterwards, we calculated the mean population activity at every time bin of the centered mean sequence. We next sought to fit a Gaussian to the bump of activity of each dataset separately using the MATLAB function “fit”.:

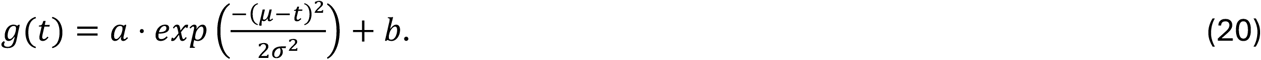

To obtain the offset *b* we identified the minimum value of the bump of activity. Then, we subtracted the offset value from the bump and fit the values *a* and *σ* of *g*(*t*) using the MATLAB function “fit”. The goodness of fit was R^2^= 0.99 for the bat data, and R^2^=0.99 for the mouse data. The parameters a, b, and σ match the amplitude, offset, and width in equation 4, which defined the rate target for each RNN unit.

#### Generation of synthetic sequences from fitted parameters

After fitting the parameters of *g*(*t*) (equation 20) for each dataset separately, we generated synthetic sequences that resembled the respective experimental data by using equation 4 with the respective number of neurons, amplitude, width, and offset, and following the procedure described in the section ‘Design of simulated target sequences’.

To compare the obtained connectivity profiles between RNNs that learned the experimental sequences and those that learned the synthetic sequences generated with the fitted parameters (Figure 3 D, F), we interpolated the connectivity profiles between distances -3 and 3 before calculating their Pearson correlation.

### Perturbation analyses

To investigate the robustness of the sequential dynamics to perturbations, and how the network activity upon perturbations was related to the connectivity profile, we perturbed the RNN activity in two different ways: through *structured* and *unstructured* perturbations, explained below.

The procedure for both types of perturbations is as follows: after training the RNNs either on synthetic or experimentally recorded sequences, we froze the weights and let the dynamics run according to equations 1-3. The obtained ‘unperturbed activity’ was 17s long for the simulated sequences (bin size = 0.0081s), 6s long for the bat sequences (bin size = 0.0081s), and 322.5s long for the rodent sequences (bin size = 0.0081s). Next, we perturbed the generated dynamics locally in time and compared the effects of the perturbations on the network activity, with the unperturbed network response.

#### Structured perturbations

We split all neurons into non-overlapping ensembles of neighbouring neurons in the sequence ordering. All ensembles had the same number of neurons, i.e., the same ‘size’. We varied the size of the ensembles such that each ensemble would have ⌊*N* ⋅ *p*_*E*_⌋ adjacent units, with *p*_*E*_ ∈ {0.1,0.2,0.3} and *N* the total number of units in the RNN. We denoted the ensembles E, …, E, where 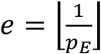 is the total number of ensembles. Next, we took the unperturbed activity and split it into two halves in the time domain. For the synthetic and rodent data, this approximately corresponded to having one sequence in the first half, and a second sequence in the second half (Figure 4A). For the bat data, this corresponded to half of the sequence in the first half, and the remaining half of the sequence in the second half. We further divided the first half of the unperturbed activity into 5 consecutive ‘time windows’, each of the same duration, denoted as T1 to T5.

As a result, we obtained a grid structure of neuronal ensembles by temporal windows. The temporal windows were 1.7s long (= 209 time bins, *dt* = 0.0081*s*) for the simulated sequences, 0.6s long (= 74 time bins, *dt* = 0.0081*s*) for the bat sequences, and 32.3s long (= 3987 time bins, *dt* = 0.0081*s*) for the mouse sequences. The size of the grid was *e* × 5. The structured perturbations consisted of perturbing the activity of one specific ensemble at one specific temporal window as described below. Outside the ensemble-temporal window pair in which the perturbation was applied, the dynamics followed equations 1-3.

Let *E* ∈ {*E*_1_, …, *E*_*e*_} be the ensemble of neurons that receives the perturbation at the temporal window *TP*.

i. Silencing the activity of one ensemble We silenced the activity of neurons in ensemble *E* at the temporal window *TP* as follows

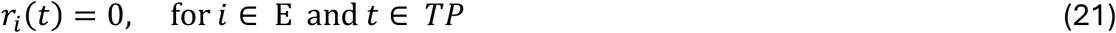

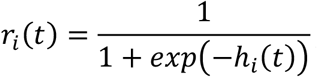otherwise.
ii. Injection of a step current into one ensemble

We injected additional current to the neurons that belong to *E* as follows

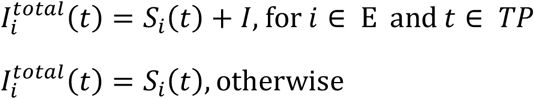

with

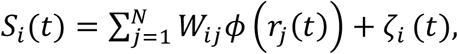

where *ζ*_*i*_(*t*) is the noise term as above (see ‘Recurrent neural network model’), and *I* taking a value from the set {0.5, 2, 5}. In Figure 4D we show an example of the breakdown of current components that add up to *I*^*total*^.

#### Unstructured perturbations

Unlike in the structured perturbations, in which we perturbed the activity of an ensemble of adjacent cells in the sequence ordering, in the unstructured perturbations we perturbed the activity of randomly selected units. We injected noise into a subset of randomly selected neurons, denoted as *Q*, during the first half of the total simulation time. We denote this as *TH*, with *TH* being equal to the time window spanned by T1 to T5 (see Figure 4A).

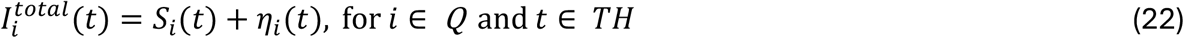

with *S*_*i*_(*t*) and *ζ*_*i*_(*t*) as above, and *η*_*i*_(*t*) is calculated as gain × *ζ*_*i*_(*t*) with *ζ*_*i*_(*t*) as in equation (2), and with gain=0.1 in Figure 4 D, gain=0.1 in Figure 4 I, gain=0.0015 in Figure 4 K, and 5, 10, 15, 20 in Supp. 9 D, E. The percentage of neurons that belonged to Q was 50% in Figure 4 C, G, H, and 10%, 25%, 50%, 75% in Supp. 9 D, E.

#### Quantification of the effect of the perturbation on the network activity

To quantify the effect of the perturbation on the network activity, we calculated the Pearson correlation between the unperturbed response and the RNN activity *after* the perturbation was released. More specifically, we defined a time window starting right after the perturbation was released until the end of the simulation. We correlated the network activity in that time window, with the unperturbed response in the analogous time window. For unstructured perturbations this entails the correlation between the unperturbed activity during the second half of the simulation, with the activity after the perturbation was released, which corresponds to the second half of the simulation with the perturbation. For structured perturbations, we obtain a correlation value for every possible combination of perturbed neural ensemble *E* ∈ {*E*_1_, …, *E*_*e*_}, and time window *TP* ∈ {*T*_1_, …, *T*_5_}. Hence, we obtain *e* × 5 correlation values that we store in a matrix *CV* of size *e* × 5 with the entry *cv*_*f,g*_ being the correlation value after perturbing ensemble *f* = 1, …, *e*, in time window *g* = 1, …,5.

Next, we quantified how the perturbation affects the dynamics as a function of how far away the perturbed ensemble was from the ensemble engaging in a sequence (see Suppl. 9). For each combination of *E* and *TP* we perturbed the activity and calculated the Pearson correlation between the perturbed and unperturbed response. Next, for each ensemble we plotted the correlation values as a function of how many time windows away *TP* was from the time window where the ensemble was engaged in the sequence. For this calculation we used periodic boundary conditions in the time windows (see for example Suppl. 9 A).

From a code perspective, each row *f* of the matrix *CV* is circularly shifted such that the entries of the diagonal in *CV* are aligned in the middle column. These entries correspond to perturbations in ensembles that were engaging during sequential activity. We calculated the mean across rows to obtain the correlation values between the unperturbed response and responses after the perturbations were released, as a function of how far away the perturbed ensemble was from the ensemble engaging in the sequence.

### Construction of upstream and downstream networks

To investigate how sequential activity in an upstream brain region generates, and constrains, sequential activity in a downstream brain region, we built two networks: an ‘upstream network’ and a ‘downstream network’.

First, we set up the upstream network as an RNN consisting of 501 neurons that was trained to generate target sequences with the specific parameters amplitude=0.5, width=1 s, offset=0.3 (bin size = 0.129 s, see ‘Recurrent neural network model’, ‘FORCE-training RNNs’, and ‘Design of simulated sequences’). We run the RNN dynamics for a total of 8.5s, which resulted in one sequence (see Figure 5B). The integration time step during training and while running the dynamics was *dt* = 0.0081 s. We used *τ* = 0.3224 s and noise *gain* = 0.5 (equations 1-3, 7-9). These parameters were kept fixed throughout all analyses in this section.

The downstream network consisted of 10 neurons that were not connected among themselves. Every neuron in the upstream network was connected to every downstream neuron. The sequential activity in the upstream network, i.e., the sequential dynamics after training the RNN, served as an input to the downstream network, such that *input*(*t*) is a column vector of size 501 × 1 that entails the response of the network at each time point *t* (bin size = 0.0081 s). The connectivity between both layers was represented by a weight matrix *A* ∈ ℝ^10×501^, where each entry *a*_*ij*_, *i* = 1, …,10, *j* = 1, . .,501 represents the weight from presynaptic upstream neuron *j* to postsynaptic downstream neuron *i*. The response of the downstream network at time *t*, i.e., the ‘output’, is denoted by the 10 × 1 vector *output*(*t*) and was calculated as the following linear readout:

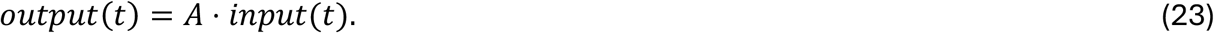

#### Design of downstream target sequences

We constructed a target activity for each downstream cell, denoted as *target*_*i*_(t), as a Gaussian that repeats over time (Equation 3, bin size = 0.0001s). Once again, we varied the triad of parameters: amplitude (a), width (σ) and offset (b), indicated below (Figure 1a). The sequence duration was 8.5 s.

Figure 5C: a=0.5, σ=2, b=0.3,

Figure 5D: a=0.5, σ=0.5, b=0.3,

Figure 5E: a=0.5, b=0.3,

Figure 5F: σ=1, b=0.3,

Figure 5G: a=0.5, σ=1,

Supp. 10 A: a=0.3, σ=0.4, b=0.3,

Supp. 10 B: a=0.5, σ=0.4, b=0.05,

Supp. 10 C: a=0.5, σ=0.4, b=0.3,

Supp. 10 D, E: a=0.5, σ=1, b=0.3.

We define *target*(*t*) = (*target*_1_(*t*), …, *target*_10_(*t*))^*T*^ as the vector of size 10 of the downstream population target activity at time *t*.

#### Training of the weights to enforce sequential activity in the downstream network

Initially, every entry in the matrix *A* was initialized at 1: *a*_*i,j*_ = 1, *i* = 1, …,10, *j* = 1, …,501. There was only one training realisation, and in that training realisation the weights in *A* were updated according to 1000 training iterations. In each training iteration *q, q* = 1, …,1000, the weights were updated once. Let

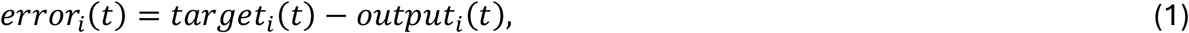

be the error of each downstream neuron’s activity over time *t* with *dt* = 0.0081, 0 ≤ *t* ≤ 8.4*s*, and *i* = 1, …,10. The weights *a*_*i,j*_ in *A* were updated as follows:

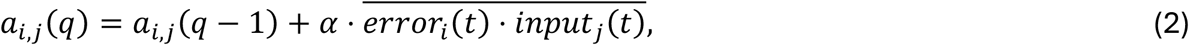

with *error*_*i*_(*t*) ⋅ *input*_*j*_(*t*) denoting the mean across time *t*, of the scalar *error*_*i*_(*t*) ⋅ *input*_*j*_(*t*). We used *α* = 0.015 as a learning rate.

We quantified the training performance of the downstream network by calculating the Pearson correlation between the matrices corresponding to the downstream population target, *target*(*t*) (see section ‘Design of downstream target sequences’), and the population output, *output*(*t*) (see section ‘Construction of upstream and downstream network’), 0 ≤ *t* ≤ 8.4 s.

### Perturbations in the upstream sequential activity

We investigated the impact of perturbations to the activity in the upstream network, on the downstream dynamics. For this analysis, the upstream and the downstream sequences both had the following properties: amplitude =0.5, width=1, and offset=0.3. After training the weights between the upstream and the downstream neurons (matrix *A*), we kept them fixed. We next introduced structured perturbations into the dynamics of the upstream network, either in the form of silencing or by additional current injection (see section ‘Perturbation analyses’). We applied the perturbation into 20% of all neurons (=100 neurons). For the injected current we used *I* = 3. Results remain unchanged when considering perturbations in 30% of all neurons. Finally, we quantified the effect of the perturbations on the downstream dynamics by calculating the correlation between the two matrices of the downstream population target, *target*(*t*) (see section ‘Design of downstream target sequences’), and the population output, *output*(*t*) (see section ‘Construction of upstream and downstream network’) during the entire simulation time 0 ≤ *t* ≤ 8.4 s.

### Software

To implement custom code, we used MATLAB (2023a).

## Notes

### Competing Interest Statement

The authors have declared no competing interest.

