## Supplementary material with analytical derivations for "A common structure in recurrent networks supports neural sequence generation locally and in downstream neurons"

### SUPPLEMENTARY MATERIAL: ANALYTICAL DERIVATIONS

This note characterizes which part of the recurrent connectivity $W$ is constrained by a target sequence. At a prescribed accuracy $\varepsilon$, the target activity and the recurrent input required to generate it are well approximated by a finite set of Fourier modes. Requiring $W$ to generate these modes fixes its action on the corresponding active subspace and, in particular, fixes the associated low-frequency Fourier components of the connectivity profile. The action of $W$ outside this active subspace remains undetermined by the sequence. We then relate this constrained component of $W$ to regularized learning and to downstream readouts.

Concretely, we show:

(1) The active Fourier modes of the connectivity profile are fixed by the target sequence (section 3).
(2) The full connectivity matrix is degenerate with respect to the target sequence and we characterize the family of matrices that generate the target sequence (section 4).
(3) Regularized learning preserves sequence-relevant Fourier modes while shrinking weakly represented modes (section 5).

#### 1. Problem formulation

We consider $N$ neurons with the standard rate model

$$\tau \dot{x} = -x + W\phi(x),$$

where $x \in \mathbb{R}^N$ is the pre-activation vector and the sigmoid $\phi(x) = 1/(1+e^{-x})$ is applied elementwise.

Let $T$ be the period of the target sequence. The target activity of neuron $n \in \{0, \dots, N-1\}$ peaks at $t = \mu_n = Tn/N$. In particular, the target activity of neuron $n$ is

$$f_n(t; a, b, \sigma) = a g_{\sigma,T}(t - \mu_n) + b,$$
$$g_{\sigma,T}(t) = \sum_{m\in\mathbb{Z}} \exp\left(-\frac{(t-mT)^2}{2\sigma^2}\right), \tag{1}$$

where $g_{\sigma,T}$ is a wrapped, i.e., periodically repeated, Gaussian. Here, $a > 0$ controls the amplitude, $b \geq 0$ the offset and $\sigma > 0$ the width of the Gaussian bump. At each time $t$, $f(t) = (f_n(t))_{n=0}^{N-1} \in \mathbb{R}^N$ is the vector of target activity across the population. We define the *target sequence* as the trajectory $t \mapsto f(t)$. Assuming $\delta < f_n(t) < 1-\delta$ for some $\delta > 0$ and all $n, t$, the *target pre-activation* is

$$x^\star(t) = \phi^{-1}(f(t)), \qquad \phi^{-1}(x) = \log(x/(1-x)).$$

**Definition 1.1.** A target sequence $f$ of period $T$ is *supported* by $W$ if

$$W f(t) = x^\star(t) + \tau \dot{x}^\star(t) := y^\star(t) \qquad \text{for all } t \in [0, T).$$

This is exactly the condition for $x^\star$ to solve the dynamics with $\phi(x^\star) = f$.

#### 2. Fourier representation and the $\varepsilon$-active sequence subspace

In the continuum limit we index neurons by $\theta \in [0, 2\pi)$, with their activity peaking at time $\mu(\theta) = T\theta/(2\pi)$, so that the population activity is a bump of activity moving along a ring of neurons with frequency $2\pi/T$:

$$f(\theta, t) = f_0(t - \mu(\theta)), \qquad f_0(t) = a\, g_{\sigma,T}(t) + b.$$

Set $\omega = 2\pi/T$ and expand $f_0(t) = \sum_{k\in\mathbb{Z}} \hat{f}_k e^{ik\omega t}$. Since $\omega\mu(\theta) = \theta$,

$$f(\theta, t) = f_0(t - \mu(\theta)) = \sum_{k\in\mathbb{Z}} \hat{f}_k e^{ik\omega t} e^{-ik\theta}.$$

The Fourier coefficients follow from $f_0$:

$$\begin{aligned} \hat{f}_k &= \frac{1}{T}\int_0^T f_0(t) e^{-ik\omega t}\,\mathrm{d}t \\ &= \frac{1}{T}\int_0^T \left(a g_{\sigma,T}(t) + b\right) e^{-ik\omega t}\,\mathrm{d}t \\ &= \begin{cases} a\hat{g}_0 + b, & k = 0, \\ a\hat{g}_k, & k \in \mathbb{Z}\setminus\{0\}. \end{cases} \end{aligned} \tag{2}$$

The Fourier coefficients of the wrapped Gaussian $g_{\sigma,T}$ are

$$\hat{g}_k = \frac{1}{T}\int_0^T g_{\sigma,T}(t) e^{-ik\omega t}\,\mathrm{d}t,$$

since $g_{\sigma,T}$ is periodic with period $T$. Substituting $g_{\sigma,T}$ with the expression in (1), we get

$$\hat{g}_k = \frac{1}{T}\int_0^T \sum_{m\in\mathbb{Z}} \exp\left(-\frac{(t - mT)^2}{2\sigma^2}\right) e^{-ik\omega t}\,\mathrm{d}t.$$

By Fubini's theorem, we may interchange the sum and the integral

$$\hat{g}_k = \frac{1}{T}\sum_{m\in\mathbb{Z}} \int_0^T \exp\left(-\frac{(t - mT)^2}{2\sigma^2}\right) e^{-ik\omega t}\,\mathrm{d}t.$$

Now make the change of variables $s = t - mT$. Since $\omega T = 2\pi$, we have

$$e^{-ik\omega t} = e^{-ik\omega(s+mT)} = e^{-ik\omega s} e^{-2\pi ikm} = e^{-ik\omega s}.$$

Therefore,

$$\hat{g}_k = \frac{1}{T}\sum_{m\in\mathbb{Z}} \int_{-mT}^{(1-m)T} \exp\left(-\frac{s^2}{2\sigma^2}\right) e^{-ik\omega s}\,\mathrm{d}s.$$

The intervals $[-mT, (1-m)T)$ tile the real line as $m$ ranges over $\mathbb{Z}$. Hence, the wrapped Gaussian unwraps to a Gaussian on $\mathbb{R}$

$$\hat{g}_k = \frac{1}{T}\int_{\mathbb{R}} \exp\left(-\frac{s^2}{2\sigma^2}\right) e^{-ik\omega s}\,\mathrm{d}s.$$

Using the Fourier transform of a Gaussian, we obtain

$$\hat{g}_k = \frac{\sigma\sqrt{2\pi}}{T} \exp\left(-\frac{1}{2}\sigma^2 (k\omega)^2\right).$$

Hence, for $k \neq 0$, the coefficient

$$\hat{f}_k = a\frac{\sigma\sqrt{2\pi}}{T} e^{-\frac{1}{2}\sigma^2 (k\omega)^2}, \tag{3}$$

decays like a Gaussian in $k$.

The target pre-activation has the same travelling-bump structure. Because

$$f(\theta, t) = f_0\,(t - \mu(\theta))\,,$$

applying the elementwise inverse nonlinearity gives

$$x^\star(\theta, t) = \phi^{-1}(f(\theta, t)) = x_0^\star(t - \mu(\theta)),$$

which is again a shifted copy of a single bump $x_0^\star(t) = \sum_k \hat{x}_k e^{ik\omega t}$. Using Definition 1.1 and differentiating in $t$ gives

$$y^\star(\theta, t) = x^\star + \tau \partial_t x^\star = \sum_{k\in\mathbb{Z}} (1 + ik\omega\tau)\hat{x}_k\, e^{ik\omega t} e^{-ik\theta}, \qquad \hat{y}_k = (1 + ik\omega\tau)\hat{x}_k.$$

Because $f$ is bounded away from the saturation of $\phi$, i.e., $f_0 \in (\delta, 1-\delta)$, the inverse $\phi^{-1}$ is smooth on the range of $f_0$. Hence, the bump $x^\star(t) = (\phi^{-1}\circ f_0)(t)$ is smooth and periodic, implying rapid decay of its Fourier coefficients $\hat{x}_k$. Since $\hat{y}_k = (1 + ik\omega\tau)\,\hat{x}_k$ it follows that $\hat{y}_k$ also exhibits rapid decay.

The wrapped Gaussian, and hence the target $f$, has nonzero Fourier coefficients at arbitrarily large $|k|$. Hence, to identify the Fourier modes relevant to the sequence at a prescribed accuracy $\varepsilon$, we introduce a cutoff using Parseval's identity. By Parseval's identity on $[0, T) \times [0, 2\pi)$,

$$\frac{1}{2\pi T}\int_0^T \int_0^{2\pi} \left| f(\theta, t) - f^{(D)}(\theta, t)\right|^2 \, \mathrm{d}\theta\, \mathrm{d}t = \sum_{|k|>D} |\hat{f}_k|^2,$$

the truncated sequence $f^{(D)}(\theta, t) = \sum_{|k|\le D} \hat{f}_k e^{ik\omega t} e^{-ik\theta}$ has mean-square error $\sum_{|k|>D} |\hat{f}_k|^2$, and likewise for $y^\star$. Since both $\hat{f}_k$ and $\hat{y}_k$ decay rapidly, for any $\varepsilon > 0$ we may define the cutoffs

$$D_f(\varepsilon) := \min\left\{ D \mid \textstyle\sum_{|k|>D} |\hat{f}_k|^2 \le \varepsilon^2 \right\},$$

and

$$D_{y^\star}(\varepsilon) := \min\left\{ D \mid \textstyle\sum_{|k|>D} |\hat{y}_k|^2 \le \varepsilon^2 \right\}.$$

We define the *sequence cutoff* as $D_{\mathsf{seq}}(\varepsilon) = \max\{D_f(\varepsilon), D_{y^\star}(\varepsilon)\}$. The associated *$\varepsilon$-active sequence subspace* is

$$\mathcal{S}_{\mathsf{seq}}(\varepsilon) := \mathrm{span}\{e^{-ik\theta} \mid |k| \le D_{\mathsf{seq}}(\varepsilon)\}.$$

That is, the space spanned by the Fourier modes needed to approximate both $f$ and $y^\star$ to an error controlled by $\varepsilon$.

*Remark* 2.1 (Finite $N$ and aliasing). A finite network samples $\theta_n = 2\pi n/N$, so the continuum mode $e^{-ik\theta}$ becomes

$$\left(v_k^{(N)}\right)_n = e^{-2\pi i k n/N}.$$

These are $N$-periodic in $k$, i.e., $v_{k+N}^{(N)} = v_k^{(N)}$. Thus, a finite population contains only $N$ distinct modes. Representing $\mathcal{S}_{\mathsf{seq}}(\varepsilon)$ without aliasing therefore requires $D_{\mathsf{seq}}(\varepsilon) < N/2$, in whcih case each active continuum mode maps to a distinct sampled mode.

Note that this means that the number of neurons required to support a sequence $f(t)$ at accuracy $\varepsilon$ is bounded below by $2D_{\mathsf{seq}}$, i.e., $N > 2D_{\mathsf{seq}}(\varepsilon)$.

#### 3. The active sequence subspace constrains the connectivity profile

In the following we write $v_k := v_k^{(N)}$ for brevity. Assume throughout that $D := D_{\mathsf{seq}}(\varepsilon) < N/2$. Equivalently, as noted in Remark 2.1, we assume that the number of neurons in the circuit is bounded below by twice the sequence cutoff $D_{\mathsf{seq}}(\varepsilon)$, i.e., $N > 2D_{\mathsf{seq}}(\varepsilon)$. Then for $|k| \leq D$ the sampled modes $(v_k)_n = e^{-2\pi i k n/N}$ are orthogonal. Let

$$\mathcal{S}_{\mathsf{seq}}^{(N)}(\varepsilon) := \mathsf{span}\{v_k \mid |k| \leq D\},$$

be the discrete $\varepsilon$-active sequence subspace. Write the $D$-mode approximations

$$f^{(D)}(t) = \sum_{|k|\leq D} \hat{f}_k e^{ik\omega t} v_k, \qquad y^{\star,(D)}(t) = \sum_{|k|\leq D} \hat{y}_k e^{ik\omega t} v_k.$$

We say $W$ *supports the active $D$-mode sequence* if $W f^{(D)}(t) = y^{\star,(D)}(t)$ for all $t \in [0, T)$.

Let $S$ be the cyclic shift, $(Sa)_n = a_{(n+1) \bmod N}$. Then, more generally,

$$(S^r a)_n = a_{(n+r) \bmod N}.$$

Since $\mu_{(n+r) \bmod N} = \mu_n + \frac{rT}{N} \bmod T$ we get

$$S^r f^{(D)}(t) = f^{(D)}\left(t - \frac{rT}{N}\right), \qquad S^r y^{\star,(D)}(t) = y^{\star,(D)}\left(t - \frac{rT}{N}\right).$$

Because $\mu_n$ are uniformly spaced on the interval $[0, T)$, shifting the neuron labels forward by $r$ positions is equivalent to shifting the sequence backward in time by $rT/N$.

The *circulant component* of $W$ is the circulant matrix given by

$$W_{\mathsf{cp}} := \frac{1}{N} \sum_{r=0}^{N-1} S^{-r} W S^r.$$

Let $d = (m - n) \bmod N$ denote the position of the presynaptic neuron relative to the postsynaptic neuron in the sequence ordering. We define the connectivity profile $\mathsf{cp} = (\mathsf{cp}_d)_{d=0}^{N-1} \in \mathbb{R}^N$ of $W$ by

$$\mathsf{cp}_d = (W_{\mathsf{cp}})_{nm}.$$

This is well defined as $W_{\mathsf{cp}}$ is circulant.

**Proposition 3.1** (The connectivity profile supports the active $D$-mode sequence). *If $W$ supports the $D$-mode sequence then it follows that*

$$W_{\mathsf{cp}} f^{(D)}(t) = y^{\star,(D)}(t),$$

*for all $t$, and in particular $W f^{(D)}(t) = W_{\mathsf{cp}} f^{(D)}(t)$.*

*Proof.* Suppose that $W$ supports the $D$-mode sequence, i.e., $W f^{(D)}(t) = y^{\star,(D)}(t)$ for all $t$. Using the shift identity

$$S^{-r} y^{\star,(D)}(t - rT/N) = y^{\star,(D)}(t),$$

it follows that

$$
\begin{aligned}
W_{\mathsf{cp}} f^{(D)}(t) &= \frac{1}{N}\sum_{r=0}^{N-1} S^{-r} W S^{r} f^{(D)}(t) \\
&= \frac{1}{N}\sum_{r=0}^{N-1} S^{-r} W f^{(D)}\left(t - \frac{rT}{N}\right) \\
&= \frac{1}{N}\sum_{r=0}^{N-1} S^{-r} y^{\star,(D)}\left(t - \frac{rT}{N}\right) \\
&= \frac{1}{N}\sum_{r=0}^{N-1} y^{\star,(D)}(t) \\
&= y^{\star,(D)}(t) \\
&= W f^{(D)}(t)
\end{aligned}
$$

where the third equality follows from the support condition. □

Thus the complementary part $W_{\perp} := W - W_{\mathsf{cp}}$ acts trivially on the $D$-mode sequence, i.e., $W_{\perp} f^{(D)}(t) = 0$ for all $t$. Since $W_{\mathsf{cp}}$ is circulant, its action on the $D$-mode sequence depends only on the Fourier modes of $\mathsf{cp}$ with $|k| \le D$. These modes are therefore fixed by the target sequence. We call the low-pass component of the connectivity profile corresponding to these modes the *$D$-mode connectivity profile* and denote it by $\mathsf{cp}^{(\le D)}$.

**Corollary 3.2** (The $D$-mode connectivity profile is fixed by the sequence)**.** *If $W$ supports the $D$-mode sequence then for every $|k| \le D$ with $\hat{f}_k \neq 0$,*

$$
W_{\mathsf{cp}} v_k = \lambda_k v_k, \qquad \lambda_k = \frac{\hat{y}_k}{\hat{f}_k} = \frac{(1 + ik\omega\tau)\hat{x}_k}{\hat{f}_k}.
$$

*Proof.* Since $W_{\mathsf{cp}}$ is circulant, the $k$th eigenvector of $W_{\mathsf{cp}}$ is the $k$th Fourier mode $v_k$. In particular, we have $W_{\mathsf{cp}} v_k = \lambda_k v_k$. Therefore,

$$
\begin{aligned}
W_{\mathsf{cp}} f^{(D)}(t) &= W_{\mathsf{cp}} \sum_{|k|\le D} \hat{f}_k e^{ik\omega t} v_k \\
&= \sum_{|k|\le D} \hat{f}_k e^{ik\omega t} W_{\mathsf{cp}} v_k \\
&= \sum_{|k|\le D} \lambda_k \hat{f}_k e^{ik\omega t} v_k.
\end{aligned}
$$

But Proposition 3.1 gives

$$
W_{\mathsf{cp}} f^{(D)}(t) = y^{\star,(D)}(t) = \sum_{|k|\le D} \hat{y}_k e^{ik\omega t} v_k.
$$

Matching the Fourier components therefore gives

$$
\lambda_k \hat{f}_k = \hat{y}_k,
$$

and hence, since $\hat{f}_k \neq 0$,

$$
\lambda_k = \frac{\hat{y}_k}{\hat{f}_k} = \frac{(1 + ik\omega\tau)\hat{x}_k}{\hat{f}_k}.
$$

□

With the convention $(W_{\mathsf{cp}})_{nm} = \mathsf{cp}_d$ with $d = (m-n) \bmod N$, the eigenvalues of the circulant matrix $W_{\mathsf{cp}}$ are given by the Discrete Fourier Transform of the connectivity profile cp. That is,

$$\lambda_k = \sum_{d=0}^{N-1} \mathsf{cp}_d e^{-2\pi i k d/N}.$$

Applying the Inverse Discrete Fourier Transform, the part of the connectivity profile fixed by the active modes with $|k| \le D$ is

$$\mathsf{cp}_d^{(\le D)} = \frac{1}{N} \sum_{|k|\le D} \lambda_k e^{2\pi i k d/N}.$$

At accuracy $\varepsilon$, the target sequence therefore fixes the low-frequency Fourier modes of the connectivity profile and leaves the remaining modes unconstrained.

#### 4. Degeneracy of the full recurrent connectivity

The $D$-mode sequence fixes the action of $W$ on $\mathcal{S}_{\mathsf{seq}}^{(N)}(\varepsilon)$. We denote its canonical extension, which vanishes on the orthogonal complement, by

$$W_D := \frac{1}{N} \sum_{|k|\le D} \lambda_k v_k v_k^* = W_{\mathsf{cp}}^{(\le D)},$$

which satisfies

$$W_D v_k = \lambda_k v_k \quad (|k| \le D). \tag{4}$$

Assuming that $\lambda_k \neq 0$ for all $|k| \le D$ it follows that $W_D$ is a sum of $2D+1$ rank-one matrices $v_k v_k^*$. Since the vectors $v_k$ are mutually orthogonal and all $\lambda_k$ are nonzero, the $2D+1$ rank-one terms act on independent one-dimensional subspaces. Therefore,

$$\mathsf{rank}(W_D) = 2D+1.$$

In other words, the frequency cutoff $D$ controls the rank of the circulant matrix $W_D$.

**Proposition 4.1** (Degeneracy of supporting matrices). *Let $P_D := \frac{1}{N} \sum_{|k|\le D} v_k v_k^*$ be the orthogonal projector onto $\mathcal{S}_{\mathsf{seq}}^{(N)}(\varepsilon)$. Then, the matrices that support the D-mode sequence are*

$$\mathcal{W}_D = \{W \mid W P_D = W_D\} = \left\{W_D + \widetilde{W} \mid \widetilde{W} P_D = 0\right\}.$$

*Equivalently, $W$ supports the D-mode sequence if and only if it agrees with $W_D$ on $\mathcal{S}_{\mathsf{seq}}^{(N)}(\varepsilon)$.*

*Proof.* First, suppose

$$W = W_D + \widetilde{W}$$

with $\widetilde{W} P_D = 0$. Since $f^{(D)}(t)$ lies in $\mathcal{S}_{\mathsf{seq}}^{(N)}(\varepsilon)$ for every $t$, we have

$$P_D f^{(D)}(t) = f^{(D)}(t).$$

Therefore,

$$\widetilde{W} f^{(D)}(t) = \widetilde{W} P_D f^{(D)}(t) = 0.$$

Hence,

$$W f^{(D)}(t) = (W_D + \widetilde{W}) f^{(D)}(t) = W_D f^{(D)}(t).$$

Using (4), we get

$$\begin{aligned} W_D f^{(D)}(t) &= W_D \sum_{|k|\le D} \hat{f}_k e^{ik\omega t} v_k \\ &= \sum_{|k|\le D} \hat{f}_k e^{ik\omega t} W_D v_k \\ &= \sum_{|k|\le D} \lambda_k \hat{f}_k e^{ik\omega t} v_k \\ &= \sum_{|k|\le D} \hat{y}_k e^{ik\omega t} v_k \\ &= y^{\star,(D)}(t). \end{aligned}$$

Thus, every matrix of the form $W_D + \widetilde{W}$ with $\widetilde{W} P_D = 0$ supports the $D$-mode sequence.

Conversely, suppose $W$ supports the $D$-mode sequence, i.e.,

$$W f^{(D)}(t) = \sum_{|k|\le D} \hat{f}_k e^{ik\omega t} W v_k = \sum_{|k|\le D} \hat{y}_k e^{ik\omega t} v_k = y^{\star,(D)}(t).$$

Hence, by uniqueness of the Fourier coefficients we have

$$\hat{f}_k W v_k = \hat{y}_k v_k, \qquad |k| \le D,$$

which implies

$$W v_k = \lambda_k v_k, \quad |k| \le D.$$

By (4) it follows that

$$(W - W_D) v_k = 0, \qquad |k| \le D.$$

Therefore,

$$(W - W_D) P_D = 0.$$

Setting $\widetilde{W} = W - W_D$ gives

$$W = W_D + \widetilde{W}, \qquad \widetilde{W} P_D = 0.$$

The claim follows. □

All matrices in $\mathcal{W}_D$ generate the same $D$-mode sequence. The free part decomposes as $\widetilde{W} = W_{\mathrm{cp}}^{(>D)} + W_{\perp}$, where $W_{\mathrm{cp}}^{(>D)}$ contains the high-frequency circulant modes and $W_{\perp}$ the non-circulant part. The former satisfies $W_{\mathrm{cp}}^{(>D)} P_D = 0$ automatically, so the constraint $\widetilde{W} P_D = 0$ amounts to requiring $W_{\perp}$ to vanish on the active subspace and is otherwise unconstrained. Thus, at accuracy $\varepsilon$, the target sequence fixes the low-pass connectivity profile but not the full matrix.

#### 5. Relation to learned connectivity

To understand how regularized learning acts on the sequence-relevant modes, we consider the idealized regression problem obtained by replacing the target sequence by its $D$-mode approximation at accuracy $\varepsilon$

$$\min_{W} \frac{1}{NT} \int_0^T \| W f^{(D)}(t) - y^{\star,(D)}(t) \|_2^2 \, \mathrm{d}t + \alpha \| W \|_F^2.$$

The first term penalizes deviations from the support condition, i.e., $W f^{(D)}(t) = y^{\star,(D)}(t)$, while the second penalizes the Frobenius norm of the connectivity.

The data are shift-equivariant, i.e., $S^r f^{(D)}(t) = f^{(D)}(t - rT/N)$ and likewise for $y^{\star,(D)}$, and the Frobenius norm $\|\cdot\|_F$ is shift-invariant. Hence, the objective is invariant under $W \mapsto S^{-r} W S^r$. Since $\alpha > 0$ makes the objective strictly convex with a unique minimizer, that minimizer must be circulant. For $\alpha > 0$, the minimizer generally trades exact support of the target sequence against a reduction in connectivity norm.

Since $D < N/2$, the retained Fourier modes are distinct in the finite population, and the optimization decouples mode by mode into

$$\min_{\lambda_k^{(\alpha)}} \left|\lambda_k^{(\alpha)} \hat{f}_k - \hat{y}_k\right|^2 + \alpha \left|\lambda_k^{(\alpha)}\right|^2, \qquad |k| \le D,$$

with ridge solution

$$\lambda_k^{(\alpha)} = \frac{\overline{\hat{f}_k}\, \hat{y}_k}{|\hat{f}_k|^2 + \alpha} = \frac{|\hat{f}_k|^2}{|\hat{f}_k|^2 + \alpha}\, \lambda_k,$$

where

$$\lambda_k = \frac{\hat{y}_k}{\hat{f}_k},$$

is the sequence-required eigenvalue derived in Corollary 3.2. Modes with $|\hat{f}_k|^2 \gg \alpha$ satisfy $\lambda_k^{(\alpha)} \approx \lambda_k$, whereas modes with $|\hat{f}_k|^2$ less than or on the scale of $\alpha$, i.e., $|\hat{f}_k|^2 \lesssim \alpha$, are strongly shrunk toward zero. Thus, regularization acts as a soft spectral cutoff. That is, modes that are strongly represented in the target sequence are fitted accurately, whereas weak high-frequency modes are increasingly sacrificed in favor of a smaller connectivity norm.

The unique optimum of this idealized regression problem is therefore a low-pass circulant approximation to the connectivity required by the target sequence. The FORCE training procedure seeks to approximate this optimum, predicting that independently trained networks share its dominant low-frequency connectivity profile. This matches our simulations.

#### 6. Parameter dependence of the learned connectivity profile

The sequence-relevant part of the learned profile is $\mathsf{cp}^{(\alpha)} = \left(\mathsf{cp}_d^{(\alpha)}\right)_{d=0}^{N-1}$ with

$$\mathsf{cp}_d^{(\alpha)} = \frac{1}{N} \sum_{|k| \le D} \lambda_k^{(\alpha)} e^{2\pi i k d/N}.$$

Hence, the sequence parameters, $a, b$ and $\sigma$, enter through

$$\lambda_k^{(\alpha)} = \frac{|\hat{f}_k|^2}{|\hat{f}_k|^2 + \alpha} \lambda_k,$$

in two ways: Through the sequence-required eigenvalue

$$\lambda_k = \frac{\hat{y}_k}{\hat{f}_k},$$

and through the shrinkage factor

$$\frac{|\hat{f}_k|^2}{|\hat{f}_k|^2 + \alpha}.$$

For $k \neq 0$, we have

$$|\hat{f}_k|^2 = a^2 \left(\frac{\sigma\sqrt{2\pi}}{T}\right)^2 e^{-\sigma^2 (k\omega)^2}.$$

Hence, increasing $a$ increases $|\hat{f}_k|^2$, which implies less shrinkage of the nonzero Fourier modes. In particular, more modes survive regularization.

Increasing $\sigma$ instead concentrates the Fourier spectrum toward low frequencies. To determine how this affects regularization, consider how $\hat{f}_k$ changes with $\sigma$:

$$\frac{\partial}{\partial\sigma}|\hat{f}_k| = a\frac{\sqrt{2\pi}}{T}e^{-\frac{1}{2}\sigma^2(k\omega)^2}\left(1-\sigma^2(k\omega)^2\right).$$

Thus,

$$\frac{\partial}{\partial\sigma}|\hat{f}_k| < 0 \qquad \text{for} \qquad |k| > \frac{1}{\sigma\omega}.$$

Hence, increasing the sequence width $\sigma$ decreases sufficiently high-frequency Fourier coefficients, causing their shrinkage factors to decrease and these modes to be more strongly suppressed by regularization. This favors flatter learned connectivity profiles.

The offset $b$ acts differently. For $k \neq 0$, it leaves $\hat{f}_k$ unchanged but shifts the required input through $x^\star = \phi^{-1}(f)$ and $y^\star = x^\star + \tau\dot{x}^\star$. Hence, the offset $b$ changes $\hat{x}_k$ and

$$\lambda_k = \frac{(1+ik\omega\tau)\hat{x}_k}{\hat{f}_k}.$$

This provides a mechanism by which the offset can alter the asymmetry of the connectivity profile.

In summary, wider sequences favor flatter learned connectivity profiles, while sequences with larger amplitudes can expose higher-frequency structure when the corresponding eigenvalues are non-negligible.

#### 7. Constraints on downstream sequences scaffolded by an upstream sequence

Let $N_u$ and $N_d$ denote the number of neurons in the upstream and downstream networks, respectively. Let

$$t \mapsto u(t) \in \mathbb{R}^{N_u}, \qquad \text{and} \qquad t \mapsto z(t) \in \mathbb{R}^{N_d},$$

denote the full upstream and downstream target sequences. For a prescribed accuracy $\varepsilon > 0$, let $u^{(\varepsilon)}$ and $z^{(\varepsilon)}$ denote their respective truncated Fourier representations, retaining all sequence-relevant modes at that accuracy. We assume that $N_u$ and $N_d$ are sufficiently large so that these modes are represented without aliasing, as in Remark 2.1.

We say that $u$ *scaffolds* $z$ at accuracy $\varepsilon$ if there exists a linear readout $J_\varepsilon \in \mathbb{R}^{N_d\times N_u}$ such that

$$z^{(\varepsilon)}(t) = J_\varepsilon u^{(\varepsilon)}(t)$$

for all $t$. We refer to *exact scaffolding* as the limit in which $\varepsilon \to 0$, the population sizes increase accordingly at a fixed ratio $N_d/N_u$, and the operator norms $\|J_\varepsilon\|_2$ remain uniformly bounded.

Let $T_u$ and $T_d$ be the fundamental periods of $u$ and $z$, respectively. Their truncated counterparts share the same fundamental periods. Suppose that $u$ scaffolds $z$ at accuracy $\varepsilon$. Since $u^{(\varepsilon)}(t+T_u) = u^{(\varepsilon)}(t)$,

$$z^{(\varepsilon)}(t+T_u) = J_\varepsilon u^{(\varepsilon)}(t+T_u) = J_\varepsilon u^{(\varepsilon)}(t) = z^{(\varepsilon)}(t).$$

Hence, $T_u$ is also a period of $z^{(\varepsilon)}$. It follows that

$$T_u = qT_d, \qquad q \in \mathbb{N},$$

or equivalently

$$T_d = \frac{T_u}{q} \leq T_u.$$

Thus, a scaffolded sequence can have a shorter fundamental period than the upstream sequence, but cannot have a longer one.

**Width.** Index the upstream neurons by $i \in \{0, \dots, N_u - 1\}$ and the downstream neurons by $j \in \{0, \dots, N_d - 1\}$. Consider Gaussian upstream and downstream sequences

$$u_i(t) = a_u g_{\sigma_u, T_u}(t - \mu_i) + b_u, \qquad z_j(t) = a_d g_{\sigma_d, T_d}(t - \nu_j) + b_d,$$

with

$$\mu_i = \frac{iT_u}{N_u}, \quad \text{and} \quad \nu_j = \frac{jT_d}{N_d}.$$

Since $T_u = qT_d$, their fundamental frequencies satisfy

$$\omega_d = \frac{2\pi}{T_d} = \frac{2\pi}{\frac{1}{q}T_u} = q\frac{2\pi}{T_u} = q\omega_u.$$

Similar to (3), for $k \neq 0$ and $m \neq 0$, the scalar Fourier coefficients are

$$\hat{u}_k = a_u \frac{\sigma_u\sqrt{2\pi}}{T_u} e^{-\frac{1}{2}\sigma_u^2(k\omega_u)^2}, \qquad \text{and} \qquad \hat{z}_m = a_d \frac{\sigma_d\sqrt{2\pi}}{T_d} e^{-\frac{1}{2}\sigma_d^2(m\omega_d)^2}. \tag{5}$$

The corresponding population Fourier components are

$$\hat{u}_k v_k^{(u)} \qquad \text{and} \qquad \hat{z}_m v_m^{(d)},$$

where $v_k^{(u)}$ and $v_m^{(d)}$ are the Fourier modes of the upstream and downstream populations, respectively.

Suppose that $u$ scaffolds $z$ at accuracy $\varepsilon$. For every retained downstream mode $m$, scaffolding requires the corresponding upstream mode $qm$ to be retained. Since

$$m\omega_d = qm\omega_u,$$

uniqueness of the Fourier coefficients in $z^{(\varepsilon)}(t) = J_\varepsilon u^{(\varepsilon)}(t)$ gives

$$\hat{z}_m v_m^{(d)} = J_\varepsilon \left(\hat{u}_{qm} v_{qm}^{(u)}\right).$$

Taking Euclidean norms gives

$$|\hat{z}_m| \|v_m^{(d)}\|_2 \leq \|J_\varepsilon\|_2 |\hat{u}_{qm}| \, \|v_{qm}^{(u)}\|_2.$$

Since

$$\|v_m^{(d)}\|_2 = \sqrt{N_d}, \qquad \text{and} \qquad \|v_{qm}^{(u)}\|_2 = \sqrt{N_u},$$

it follows that

$$\|J_\varepsilon\|_2 \geq \sqrt{\frac{N_d}{N_u}} \frac{|\hat{z}_m|}{|\hat{u}_{qm}|}.$$

Substituting the Gaussian Fourier coefficients from (5) and using $T_u = qT_d$ and $\omega_d = q\omega_u$ gives

$$\|J_\varepsilon\|_2 \geq q\sqrt{\frac{N_d}{N_u}} \frac{a_d\sigma_d}{a_u\sigma_u} \exp\left[\frac{1}{2}(qm\omega_u)^2(\sigma_u^2 - \sigma_d^2)\right]. \tag{6}$$

Note that (6) must hold for *all* modes $m$ with $1 \leq |m| \leq D_d(\varepsilon)$, where $D_d(\varepsilon)$ is the downstream Fourier cutoff at accuracy $\varepsilon$. Therefore, if the downstream sequence is narrower than the upstream sequence, i.e., $\sigma_d < \sigma_u$, then $\sigma_u^2 - \sigma_d^2 > 0$, and the right-hand side of (6) increases with $|m|$. The strongest lower bound is therefore obtained at $|m| = D_d(\varepsilon)$. That is,

$$\|J_\varepsilon\|_2 \geq q\sqrt{\frac{N_d}{N_u}} \frac{a_d\sigma_d}{a_u\sigma_u} \exp\left[\frac{1}{2}(qD_d(\varepsilon)\omega_u)^2(\sigma_u^2 - \sigma_d^2)\right]. \tag{7}$$

At finite accuracy $\varepsilon$, (7) gives a finite lower bound on $\|J_\varepsilon\|_2$. As $\varepsilon$ decreases, the downstream cutoff $D_d(\varepsilon)$ increases, since more Fourier modes must be retained to achive the prescribed accuracy. Thus, a narrower downstream sequence is not excluded at finite accuracy, but requires an increasingly large operator norm as higher-frequency modes must be retained.

For exact scaffolding, $\varepsilon \to 0$. Since the Fourier coefficients of the Gaussian sequences are nonzero at arbitrarily large $|m|$, we have

$$D_d(\varepsilon) \xrightarrow{\varepsilon \to 0} \infty.$$

Thus, if $\sigma_d < \sigma_u$, the lower bound in (7) diverges. Hence, the operator norms $\|J_\varepsilon\|_2$ cannot remain uniformly bounded as $\varepsilon$ tends to 0. Thus, exact linear scaffolding requires

$$\sigma_d \geq \sigma_u.$$

In other words, exact linear scaffolding rules out downstream Gaussian sequences that are narrower than the upstream sequence. This is consistent with our simulations.

**Amplitude, offset, and perturbations.** Once the required temporal modes are present, $J$ can rescale them, so amplitude and offset are less constrained than period and width. Perturbations propagate directly through the readout. Indeed, if $z(t) = Ju(t)$ and $u \mapsto u + \delta u$, then $\delta z(t) = J\delta u(t)$, so disruptions of the upstream sequence produce downstream errors through the scaffold itself.

Finally, by section 3, the active modes of the upstream sequence also determine the corresponding Fourier modes of its connectivity profile. These modes determine which temporal modes are available to a downstream linear readout, linking the connectivity profile associated with the upstream sequence to the family of sequences that can be scaffolded.
